# Short-hairpin RNA-guided single gene knockdown reverses triple-negative breast cancer

**DOI:** 10.1101/418764

**Authors:** Dong-Mei Chen, Zhu-Qing Shi, Li-Li Tan, Yan-Ping Chen, Chang-Qing Li, Qi Wang, He Li, Meng-Le Zhang, Jian-Ping Song, Qin Xu, Qing-Ping Zeng

## Abstract

**Background:** The origin of breast cancer remains poorly understood. Here, we testify a putative mechanism of “breast cancer origin from inducible nitric oxide synthase (iNOS) activation and oestrogen receptor alpha (ERα) inactivation”, which are classified as the essential outcomes of chronic inflammatory responses.

**Methods:** To reverse breast cancer status, iNOS was downregulated by short-hairpin RNA (shRNA)-guided *NOS2* knockdown from human triple-negative breast cancer (TNBC) cells. To re-enact breast cancer origin, ERα was downregulated by shRNA-directed *ESR1* knockdown in human mammary epithelial cells.

**Results:** Upon *NOS2* knockdown from HCC1937 cells, the specific TNBC transcription factor genes, *RUNX1* and *BCL11A*, were downregulated, hypoxia was compromised, Warburg effects were attenuated, and tumourigenic proliferation was halted, accompanied by an increase in the tumour marker cyclin D1 (CD1) and a decrease in the tumour suppressor cyclin-dependent kinase inhibitor (CKI). In contrast, *ESR1* knockdown from MCF-10A cells led to upregulated *BCL11A* and *RUNX1* expression, augmented hypoxic responses, pronounced Warburg effects, and enhanced PI3K/Akt/mTOR signalling, together with low levels of CD1 and high levels of CKI induction.

**Conclusions:** Breast cancer should originate from inflammatory signalling, during which iNOS activation and ERα inactivation elicit hypoxia, oxidation, mitochondrial dysfunction, and breast cancer-like hyperplasia, demonstrating that iNOS inhibitors and ERα activators represent promising candidate prodrugs enabling breast cancer prevention in the early stage.

## Background

Breast cancer affects approximately 12% of women worldwide and affected 2.8 million American women in 2015 [1]. Hormone receptor-positive (HR+) breast cancer constitutes more than 70% of patients among postmenopausal women [2, 3]. The incidence of HR+ breast cancer has been anticipated to increase by 4% annually between 2011 and 2030 in women aged 70-84 years old and 1.6% among women aged 50-69 years old [4]. While 2-3% of the family-transmitted and early-onset patients can be attributed to the germ-line mutation genes *BRCA1* and *BRCA2* [5], most sporadic and late-onset clinical cases originate from versatile risk factors.

Although the aetiological initiators of breast cancer remain to be identified, the female hormones oestrogens, mainly including estrone (E1), estradiol (E2) and estriol (E3), have been postulated to promote mammary tumourigenesis via activation of oestrogen receptor (ER)-induced downstream pro-mitogenic transcriptional mechanics [6]. The ER-independent effects of oestrogens have been shown to also influence breast tumour development in concert with ER-dependent effects [7]. Recently, evidence from clinical and experimental data has emerged to link the gut microbiota to breast cancer [8]. This relevance was built on 40 years of literature regarding excreted oestrogen recycling via deconjugation by β-glucuronidase (GUS) from gut commensal microbes [9]. Intestinal microbial richness and enzymatic activity, including but not limited to GUS, have been shown to influence the levels of non-ovarian oestrogens through the enterohepatic circulation [10]. Indeed, high levels of oestrogens were observed to correlate to a high incidence of breast cancer [11]. Women with a more diverse gut microbiome have been shown to exhibit elevated urinary hydroxylated oestrogen metabolites [12].

During ER-independent mammary tumourigenesis, oestrogens are converted to 2-hydroxyestradiol (2-OHE2), 4-hydroxyestradiol (4-OHE2), semi-quinones, and quinones by cytochrome P450 enzymes, among which quinones can react with DNA to produce apurinic nucleotide and lead to A-T to G-C transition [13]. The highly depurinated adducts have also been detected in serum and urine samples from breast cancer patients and women with a strong family history of breast cancer [14]. Mouse gene mutations induced by 2-OHE2 and 4-OHE2 have been confirmed to cause uterine cancer [15] and kidney cancer [16]. The concentration of 4-OHE2 in breast cancer biopsies was found to be 3 times higher than in normal breast tissue [17], and urinary 2-OHE2 and 4-OHE2 levels were elevated in breast cancer patients compared with healthy controls [18].

Regarding the ER-dependent mammary tumourigenic mechanism, it remains unclear how can inflammation induce mammary hyperplasia. According to clinical data indicating a family-transmitted ERα mutation (R394H) with hyperoestrogenemia (>50-fold normal range) [19], we anticipated that inactivation of *ESR1*-encoding ERα would result in oestrogen insensitivity and oestrogenemia. Because pro-inflammatory cytokines can activate *NOS2*-encoding inducible nitric oxide synthase (iNOS) for nitric oxide (NO) generation and peroxynitrite (ONOO^-^) formation, NO- and ONOO^-^-triggered ERα nitrosylation can affect ERα activity [20]. Therefore, iNOS activation and ERα inactivation should induce oestrogenemia and provoke mammary tumourigenesis or carcinogenesis.

Based on the above reasoning, we suggest herein a hypothesis of “breast cancer origin from iNOS activation and ERα inactivation” for deciphering ER-dependent mammary tumourigenesis. To verify this hypothesis, we intend to silence *NOS2* and prohibit iNOS activation for reversing tumour progression, on the one hand, and to silence *ESR1* and mimic ERα inactivation for imitating tumour origin, on the other hand. Until now, no loss-of-function tests of *NOS2* and *ESR1* in tumour and normal cells, respectively, have been conducted to testify this putative tumour-inducing possibility.

Triple-negative breast cancer (TNBC), referring to any breast cancer that does not express ER, progesterone receptor (PR), and human epidermal growth factor 2 (HER2) receptor, is associated with a poor prognosis [21]. TNBC represents the female hormone analogue therapy-intractable subtype of breast cancer, accounting for 15% of clinical cases [22]. Using multivariate analysis, the new prognostic breast cancer marker *RUNX1* was identified to correlate with the poor prognosis specifically in TNBC [23]. By analysing the expressional profiles, it was found that *BCLllA* is highly expressed in TNBC, and its overexpression in non-tumourigenic cells drives tumour development. In contrast, *BCL11A* downregulation in TNBC cell lines leads to a reduction of tumour size [24].

To reverse or re-enact the expression and metabolic profiles of breast cancer, *NOS2* or *ESR1* was downregulated in the human TNBC cell line HCC1937 or the human mammary epithelial cell line MCF-10A. Additionally, E2 was supplemented in the latter to create a milieu mimicking the dysfunctional *ESR1*-induced oestrogenemia. We demonstrated for the first time that *NOS2* knockdown could convert breast cancer cells to nearly normal cells, whereas *ESR1* knockdown could transform normal cells to breast cancer-like cells. This work, with prospects for an effective breast cancer therapy through the use of iNOS inhibitors and ERα activators, should shed light on the exploration of breast cancer origin, extend our scope of understanding regarding the mechanism underlying mammary tumourigenesis/carcinogenesis, and provide attractive and effective treatment regimens in hormone replacement-intractable TNBC patients

## Methods

### Cell culture, staining and counting

The TNBC cell line HCC1937 (ATCC) was cultured in 1640 medium (Gibco) supplemented with 10% foetal bovine serum (FBS, Gibco). The mammary epithelial cell line MCF-10A (ATCC) over-expressing the recombinant *ESR1* gene (LangRi, Guangzhou, China) was cultured in DMEM/F12 medium (Gibco) supplemented with 10% FBS (Gibco). Cell thawing and subculture were conducted according to the standard procedures provided by the manufacturers. To plot the growth curve, live cells were stained with 3-(4,5-dimethyl-2-thiazolyl)-2,5-diphenyl-2-H-tetrazolium bromide (MTT), and the absorbance was measured at 490 nm. To count the colony numbers, cells in the exponential growth phase were diluted to 500 cells/ml, cultured for 7-10 d, and stained with haematoxylin.

### Vector construction and stable transfection

The retrovirus-based and integration-type plasmid expression vector pSuper-retro-puro (OligoEngine) was inserted separately into one of the *NOS2-* or *ESR1*-targeted shRNA fragments, siN1/2/3 or siE1/2/3, amplified using the primer pairs listed in Table 1 to obtain the recombinant constructs psiN1/2/3 or psiE1/2/3. After transient transfection and interference with efficiency identification, psiN1/2/3 or psiE1/2/3 was exploited to transfect HCC1937 or MCF-10A for the establishment of a stable cell line. The procedures for restrictive digestion, ligation, transformation, and transfection were performed according to manufacturers’ manuals.

### knockdown vector construction and transient expression

**Table 1.**
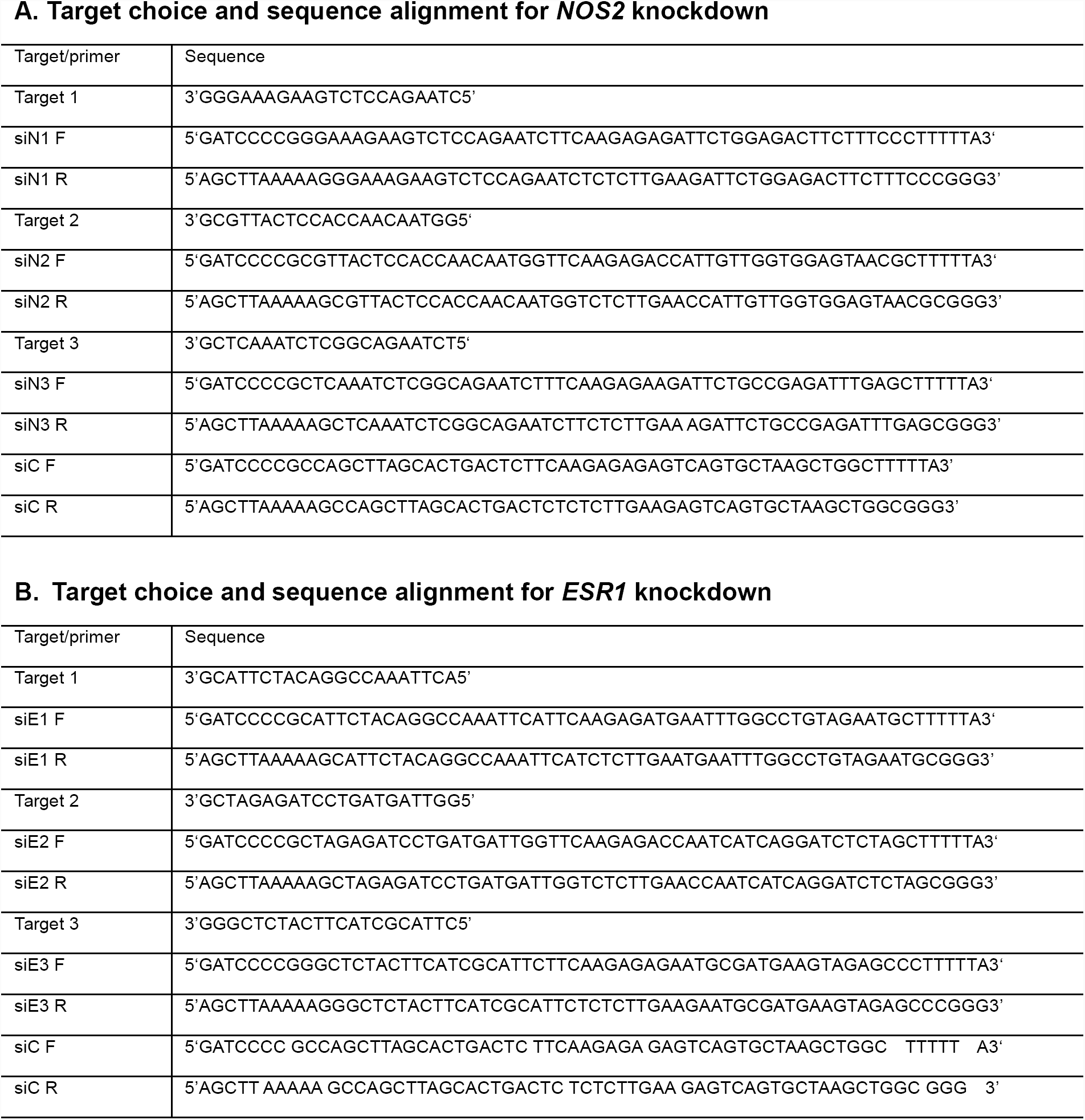
Target sequence selection and primer design for *NOS2/ESR1*.

**Table 2.**
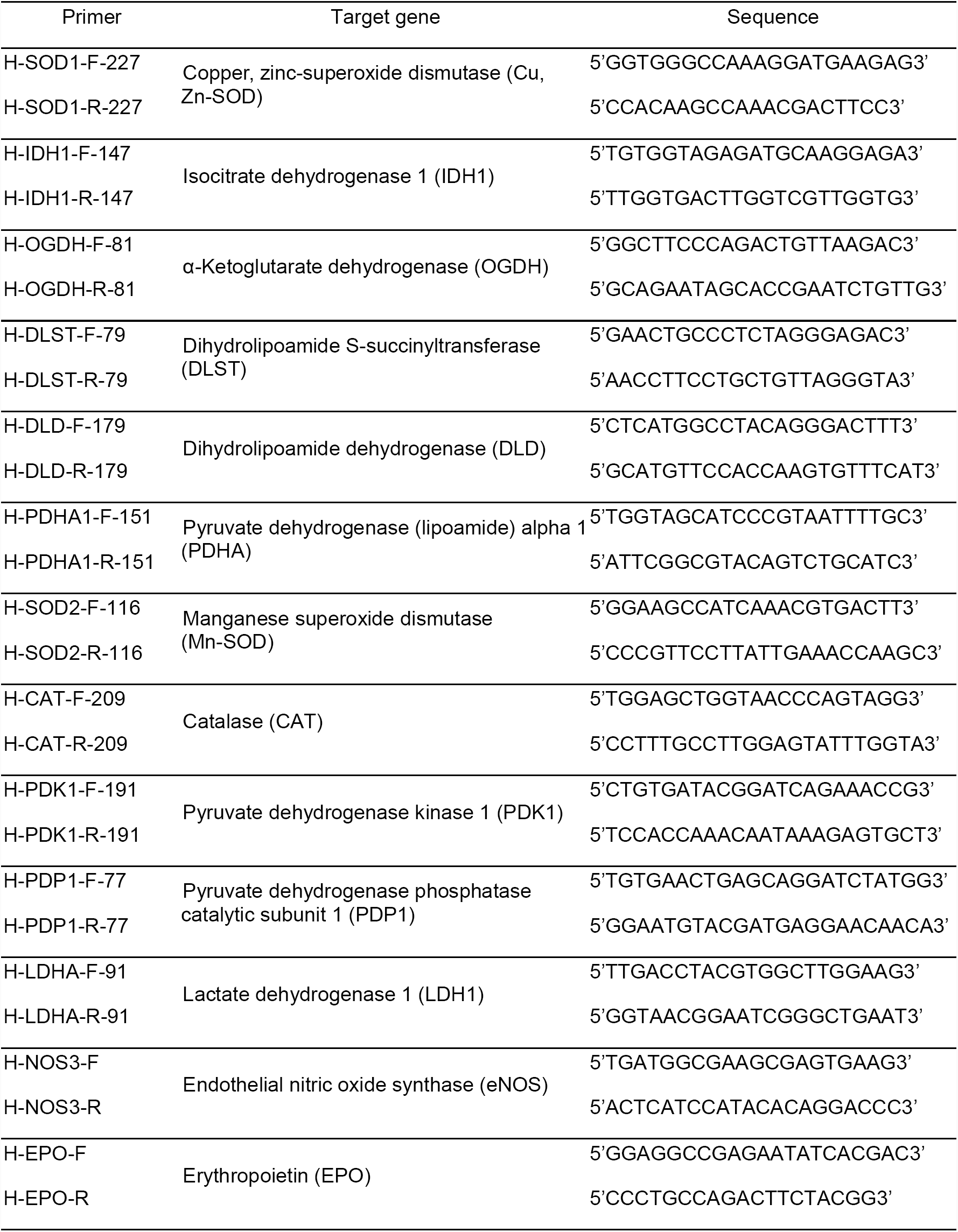
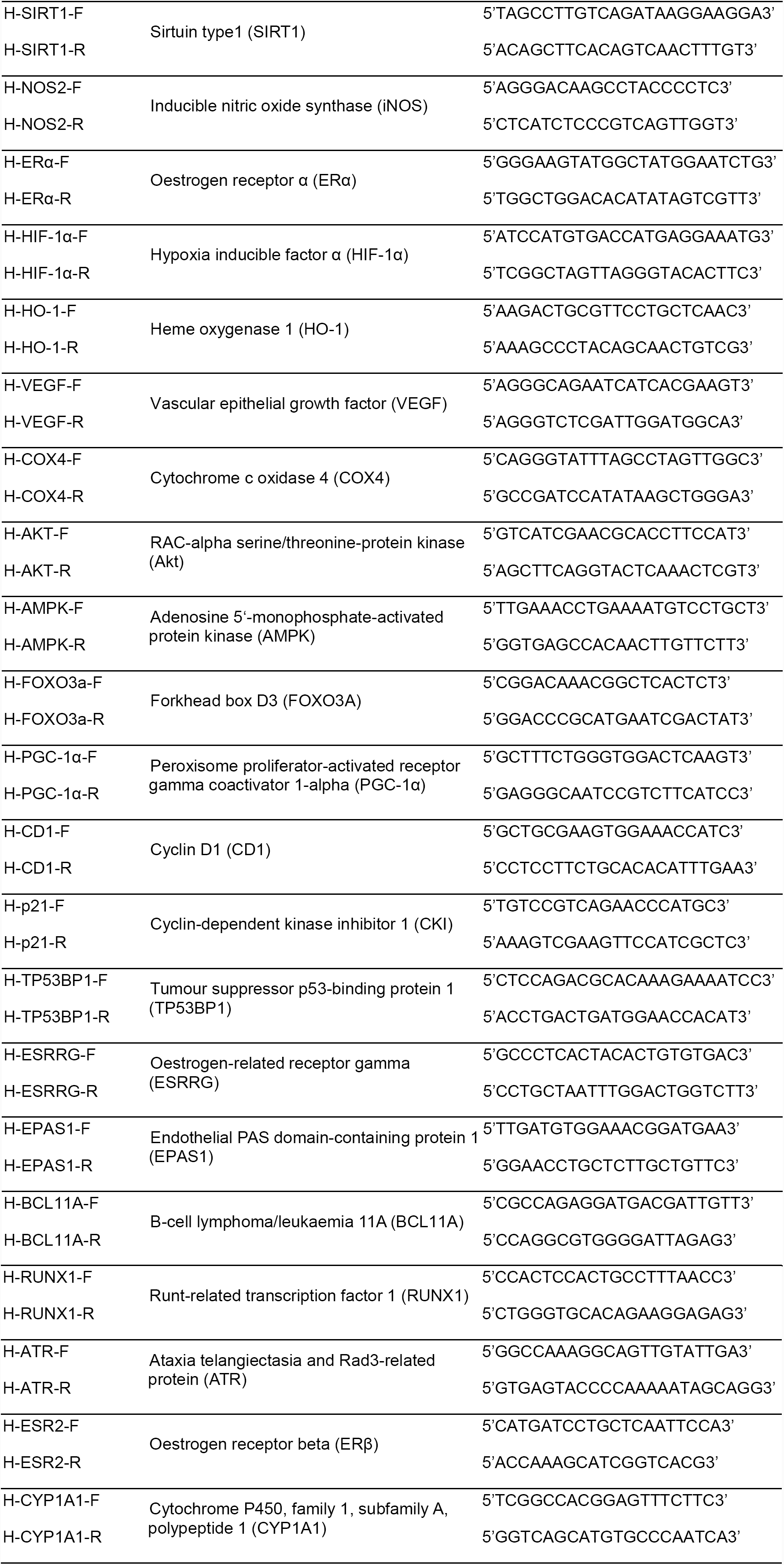
Primer sequences for target gene amplification.

### Tumour cell grafting on nude mice

BALB/c nude mice (5 weeks old) purchased from SKA, Changsha, China were housed with AL chow and free access to water drinking. For grafting, tumour cells were digested with trypsin, followed by centrifugation, re-suspension, and dilution to 1^x^ 10^5^ cells/ml. The cell suspension (100 ¼l) was subcutaneously injected into the hind legs of nude mice.

### Quantitative polymerase chain reaction (qPCR) and reverse transcription (RT)-PCR microarray

RNA isolation, purity determination, electrophoresis monitoring, reverse transcription, and quantification were performed according to the standard protocol. The relative copy numbers = 2^-Δ ΔCT^, where ΔCT = Ct target gene - Ct reference gene and Δ ΔCT = ΔC treatment sample - ΔC control sample. The primers listed in Table 2 were designed and applied under the following amplification condition: 95°C, 60 s; and 95°C, 30 s, 60°C, 35 s for 40 cycles. The RT^2^ Profiler^™^ PCR Array Mouse Fatty Liver (PAMM-157Z) was purchased from SABioscience Qiagen, Hilden, Germany. The experiment was performed by Kangchen, Shanghai, China.

### Western blotting (WB) and enzyme-linked immunosorbent assay (ELISA)

The reference protein glyceraldehyde-3-phosphate dehydrogenase (GAPDH) and antigen proteins were immunoquantified by WB according to the manufacturers’ manuals. Antibodies, including anti-PGC-1α, anti-VEGF, anti-CD1, anti-AMPKα1+AMPKα2, anti-AMPKα1 (phospho T183)+AMPKα2 (phospho T172), anti-pan-Akt, anti-pan-Akt (phospho T308), anti-eNOS, anti-iNOS, anti-α-tubulin, anti-HIF-1α, anti-HO-1, anti-EPO, anti-COX4, anti-CKI/p21, anti-SIRT1, anti-FOXO3A, anti-FOXO3A (phospho S253), and anti-Ki67 (PCNA), were purchased from Abcam (Shanghai, China). Anti-E-cadherin was provided by BOSTER (Wuhan, China), and anti-mTOR was provided by CST (Shanghai, China).

### Histochemical and immunohistochemical analysis

Formalin fixation, paraffin embedding and deparaffinization were performed as described for the haematoxylin and eosin (HE) staining procedure. Sections were incubated at room temperature with 3% H_2_O_2_ to block endogenous peroxidase and then repaired in boiling citric acid. After washing in phosphate-buffered solution, the sections were blocked with 2% bovine serum albumin and incubated with the 1:100 diluted primary antibodies at 37°C for 1 h. The primary antibody against 3-nitrotyrosine (NT) was purchased from Abcam. After washing, the sections were incubated with biotinylated secondary antibodies at 37°C for 20 min. After washing again, the sections were incubated with diaminobenzidine for 1-5 min. After rinsing with tap water, the sections were counter-stained with haematoxylin. After completion of dehydration, clearance and mounting, images were obtained under an OLYMPUS BX-51 microscope.

### Electronic microscopy

After treatment, the cells were harvested and fixed in 2.5% glutaraldehyde in 0.1 M phosphate buffer for three hours at 4°C, followed by post-fixation in 1% osmium tetroxide for one hour. Samples were dehydrated in a graded series of ethanol baths and infiltrated and embedded in Spurr’s low-viscosity medium. Ultra-thin sections of 60 nM were cut in a Leica microtome, double-stained with uranyl acetate and lead acetate, and examined using a Hitachi 7700 transmission electron microscope at an accelerating voltage of 60 kV.

### Apoptosis and reactive oxygen species (ROS) detection

The Annexin V-fluorescein isothiocyanate (FITC) Apoptosis Detection Kit was purchased from Sigma-Aldrich (USA). Cells were suspended in binding buffer and mixed with Annexin V-FITC and propidium iodide (PI) for the cytometric assay. The FITC-labelled TdT-mediated dUTP nick-end labelling (TUNEL) apoptosis detection kit was purchased from Keygen (Nanjing, China). Cells were incubated with the TdT enzyme reaction solution for fluorescence microscopic analysis. The ROS detection kit was purchased from Beyotime (Guangzhou, China). Cells were incubated with the diluted dichlorodihydrofluorescein diacetate (DCFH-DA) for detection using the fluorescence microplate reader at a wave length of 488/525 nm.

### Chorioallantoic membrane tests

The one-week-old embryonated egg was drilled to place a piece of sterilised filter paper on the 0.5-cm-diameter chorioallantoic membrane surface, and then it was loaded with 200 μl of the conditional culture medium suspension containing HCC1937::psiC or HCC1937::psiN3 cells. After sealing and incubation at 37°C with a humidity of 80% for 48-72 h, the chorioallantoic membrane was sheared to obtain images.

### Spectrophotometry

The lactic acid (LA) and adenosine triphosphate (ATP) reagent kits were purchased from Jiancheng Biotech, Nanjing, China. All determination procedures were conducted according to the manufacturers’ instructions.

### Statistical analysis

The software SPSS 22.0 was employed to analyse the raw data, and the software GraphPad Prism 5.0 was employed to plot the graphs. The independent sample test was used to compare all groups, and the Kruskal-Wallis test followed by the Nemenyi test was used when the data distribution was skewed. The significance level (p value) was set at <0.05 (*), <0.01 (**), <0.001 (***) and <0.0001 (****).

## Results

### *NOS2* knockdown results in a decline in iNOS and expression level changes of oncogenes, tumour suppressor genes and TNBC-specific transcription factor genes

Three recombinant *NOS2*-targeted small interfering RNA (siRNA) expression constructs, psiRNA-NOS1/2/3 (abbr. psiN1/2/3), were available by inserting one of three candidate shRNA fragments (see the sequence alignment in Methods), *NOS2* shRNA1/2/3, into the integration-type plasmid expression vector pSUPER-retro-puro. After separately transferring these vehicles into HCC1937 cells, *NOS2* mRNA and iNOS levels were quantified in HCC1937::psiN1/2/3 cells transiently expressing *NOS2* shRNA.

Compared with the control HCC1937::psiC cells, *NOS2* mRNAs and iNOS levels declined to different extents in HCC1937::psiN1/2/3 cells (Fig. 1a and 1b). The expression profiles showed that psiN3 exhibited a high efficiency for interfering with *NOS2* expression among others, so psiN3 was chosen to stably transfect HCC1937 cells to gain HCC1937::psiN3 cells with permanent *NOS2* shRNA expression.

**Fig. 1.**
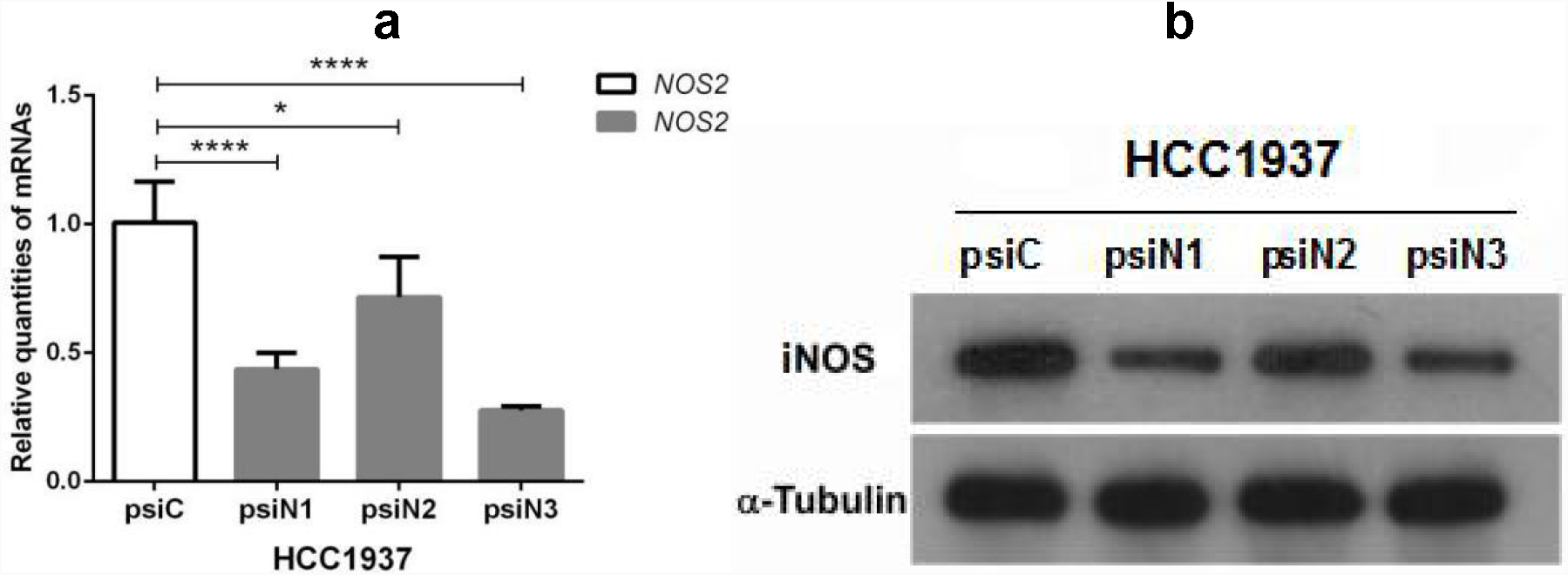
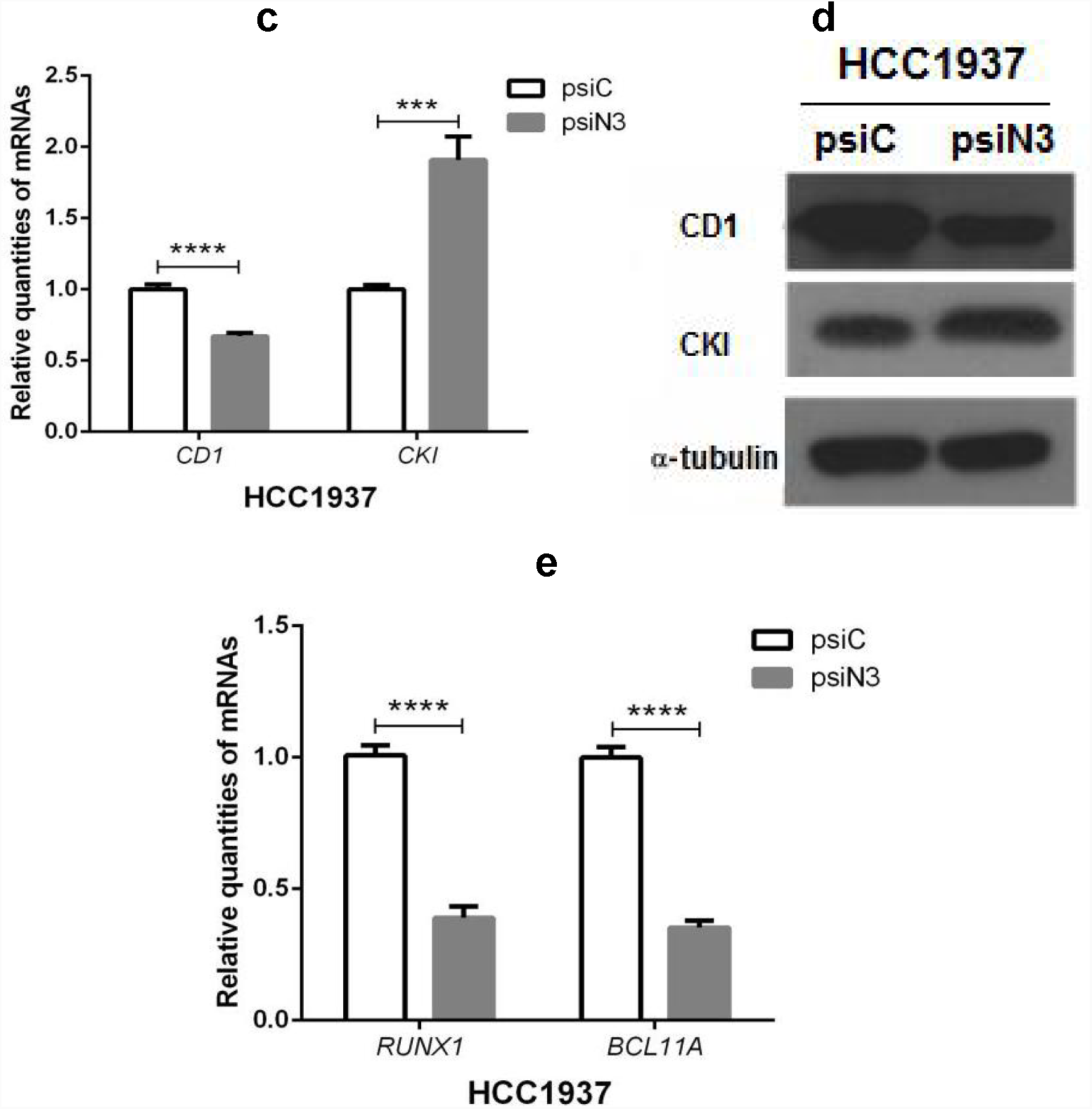
The expression profiles of *NOS2* mRNA/iNOS, *CD1* mRNA/CD1, *CKI* mRNA/CKI, *RUNX1* mRNAand *BCL11A* mRNA after shRNA-guided *NOS2* knockdown in HCC1937::psiC/psiN1/2/3 cells. **a** *NOS2* mRNA levels (transient expression); **b** iNOS levels (transient expression); **c** *CD1* and *CKI* mRNA levels; **d** CD1 and CKI levels; **e** *RUNX1* and *BCL11A* mRNA levels.

By monitoring the dynamic changes in tumourigenesis-related markers upon targeted gene knockdown, we observed that the expression levels of tumour marker CD1-encoded mRNA levels declined, whereas tumour suppressor CKI/P21-encoded mRNA levels were elevated in HCC1937::psiN3 cells (Fig. 1c). Similarly, while CD1 levels declined, CKI levels exhibited an elevation in HCC1937::psiN3 cells (Fig. 1d).

By quantifying the TNBC-specific transcription factors *RUNX1* and *BCL11A* mRNA levels, we noticed that *NOS2* knockdown could markedly downregulate *RUNX1* and *BCL11A* expression (P<0.0001) (Fig. 1e), suggesting that *NOS2* activated by pro-inflammatory cytokines might represent a pivotal step toward breast cancer.

### *NOS2* **downregulation renders** *NOS3* **upregulation with a high level of NO but a low level of ROS that unalters nitrosylation and apoptosis**

To assess the impact of *NOS2* knockdown on the correlation of *NOS2/* iNOS with *NOS3*/eNOS, we quantified the *NOS2* and *NOS3* mRNA levels (Fig. 2a) as well as the iNOS/eNOS levels (Fig. 2b) in HCC1937::psiC/psiN3 cells. As noted, the decline in *NOS2* mRNA and iNOS levels was accompanied by an elevation of *NOS3* mRNA and eNOS levels. Accordingly, an elevated NO level (Fig. 2c) was correlated with a decline in the ROS level (Fig. 2d). At a low level of ROS, ROS-scavenging SOD and CAT levels both declined (Fig. 2e and 2f).

**Fig. 2.**
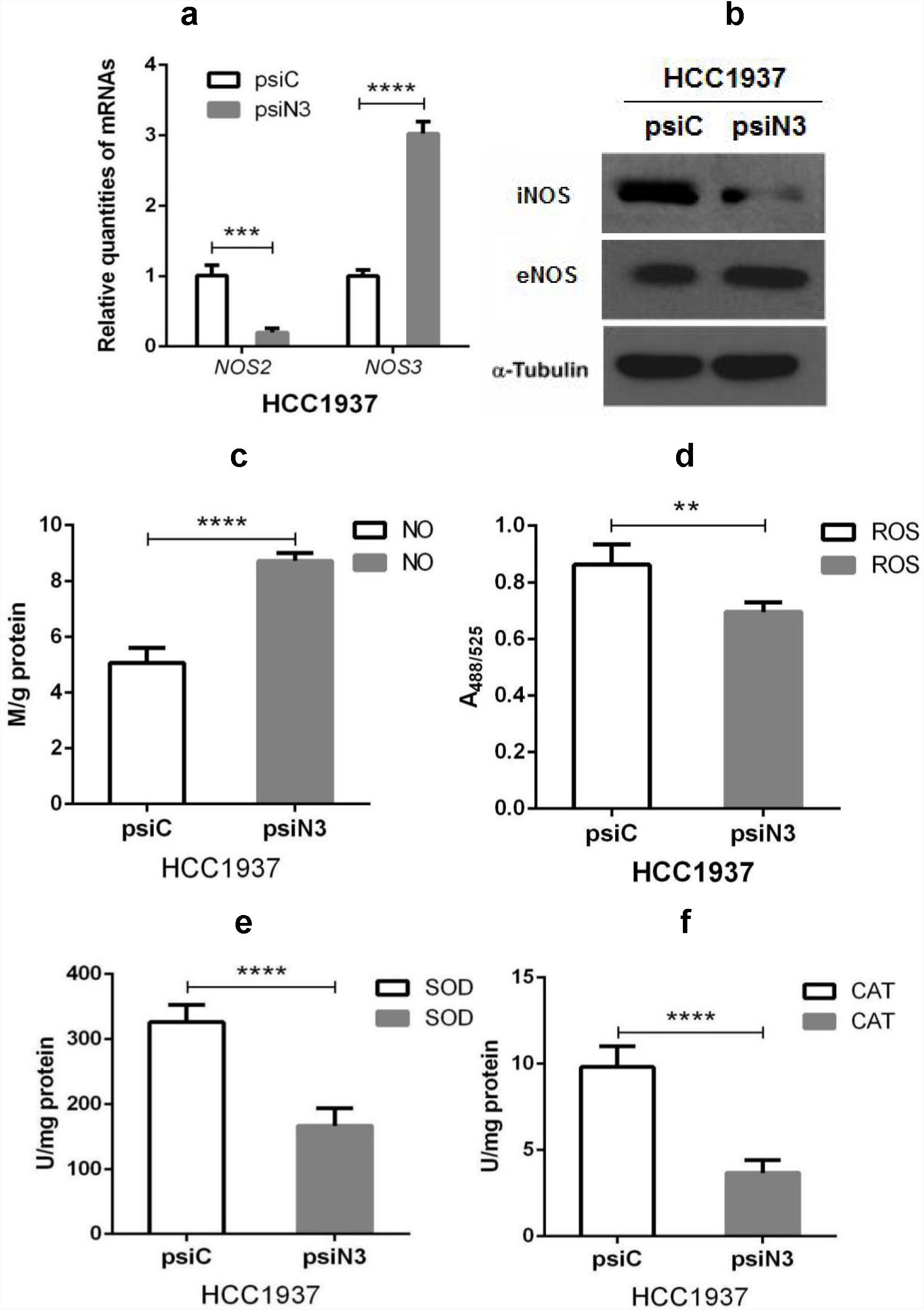
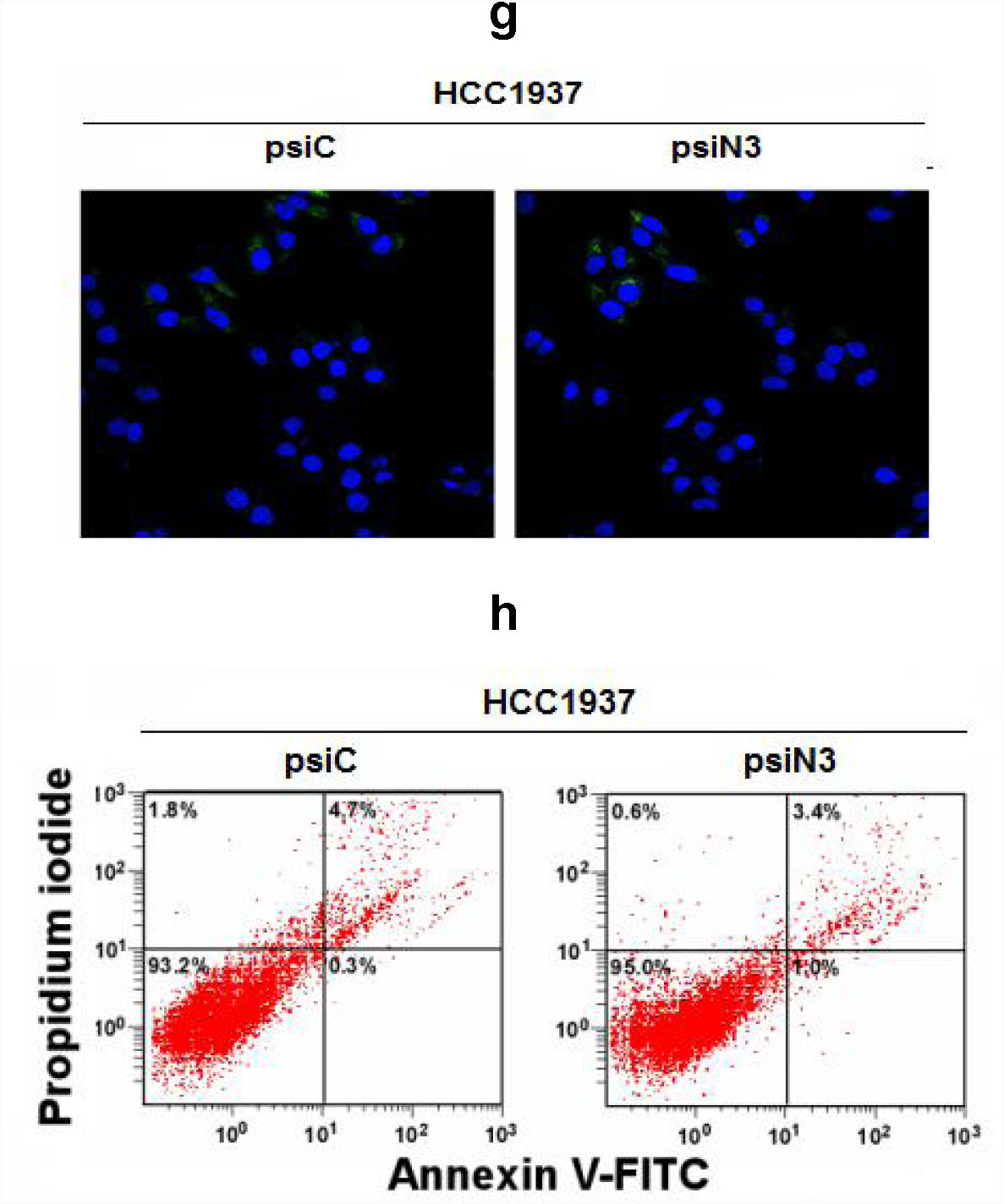
Effects of *NOS2* knockdown on *NOS2* mRNA/iNOS, *NOS3* mRNA/eNOS, NO, ROS, SOD, CAT levels, nitrosylation and apoptosis in HCC1937::psiC/psiN3 cells. **a** *NOS2* and *NOS3* mRNA levels; **b** iNOS and eNOS levels; **c** NO levels; **d** ROS levels. **e** SOD levels; **f** CAT levels; **g** NT levels in HCC1937::psiC cells (left) and HCC1937::psiN3 cells (right); **h** apoptosis percentages in HCC1937::psiC cells (left) and HCC1937::psiN3 cells (right).

Because of low-level ROS, the ROS-NO reaction-derived product ONOO^-^ should decrease accordingly. Therefore, the green fluorescence-labelled proteins carrying the ONOO^-^-modifying tyrosine residue 3-nitrotyrosine (NT) were almost equivalent between HCC1937::psiC and HCC1937::psiN3 cells (Fig. 2g). As an essential outcome, the early-phase and late-phase apoptotic cell percentages were also identical compared HCC1937::psiC cells with HCC1937::psiN3 cells (5.0% versus 4.4%) (Fig. 2h).

### An attenuated Warburg effect correlates with accelerated glucose transport into mitochondria for Krebs cycle reactions

To evaluate whether *NOS2* knockdown would affect the metabolic patterns of tumour cells, we determined the levels of ATP, an aerobic glucose catabolic end product, and LA, an anaerobic glucose catabolic end product, in HcCl937::psiC/psiN3 cells. As a result, while ATP increased (Fig. 3a), LA decreased (Fig. 3b), suggesting a metabolic conversion from glycolysis to Krebs cycle/citric acid cycle reactions, i.e., the Warburg effect was attenuated upon *NOS2* knockdown in TNBC cells.

**Fig. 3.**
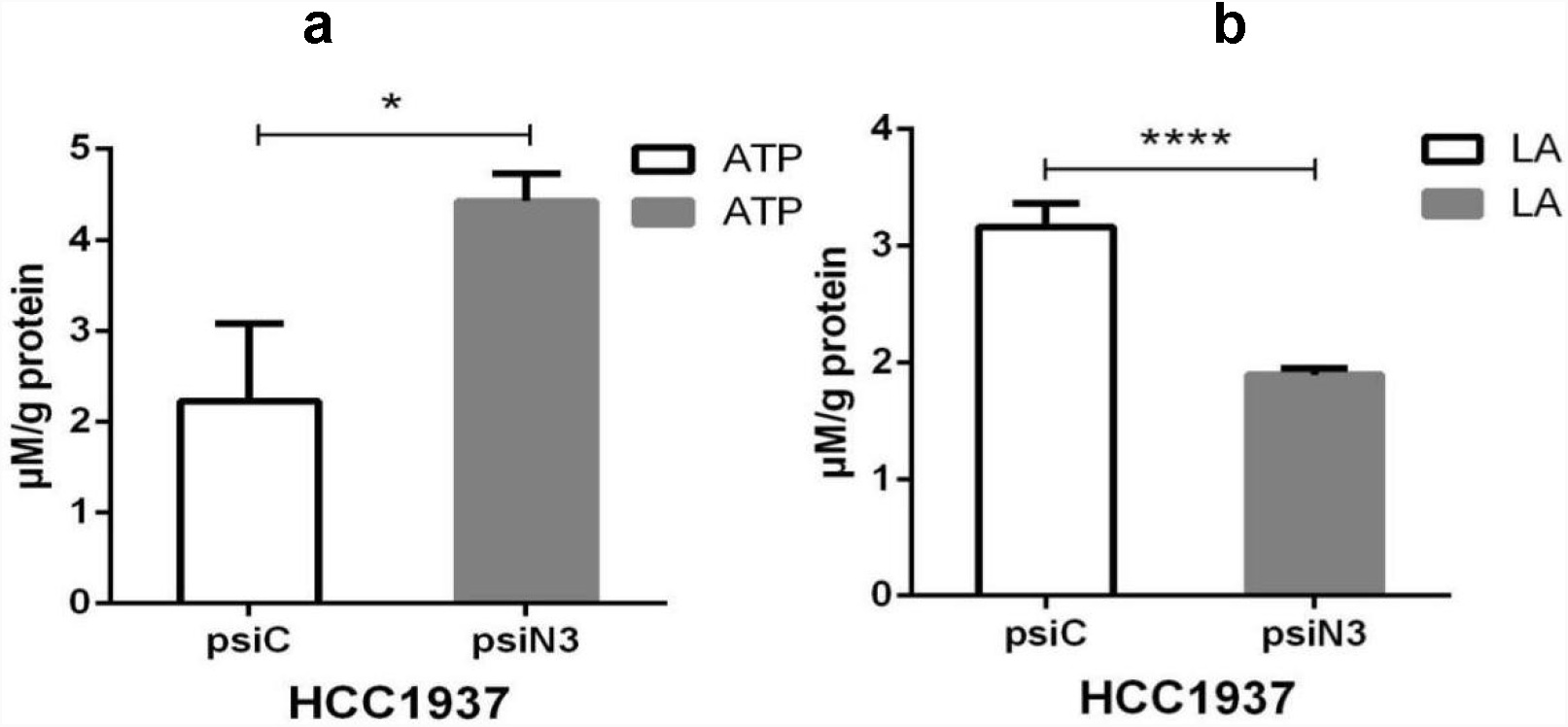
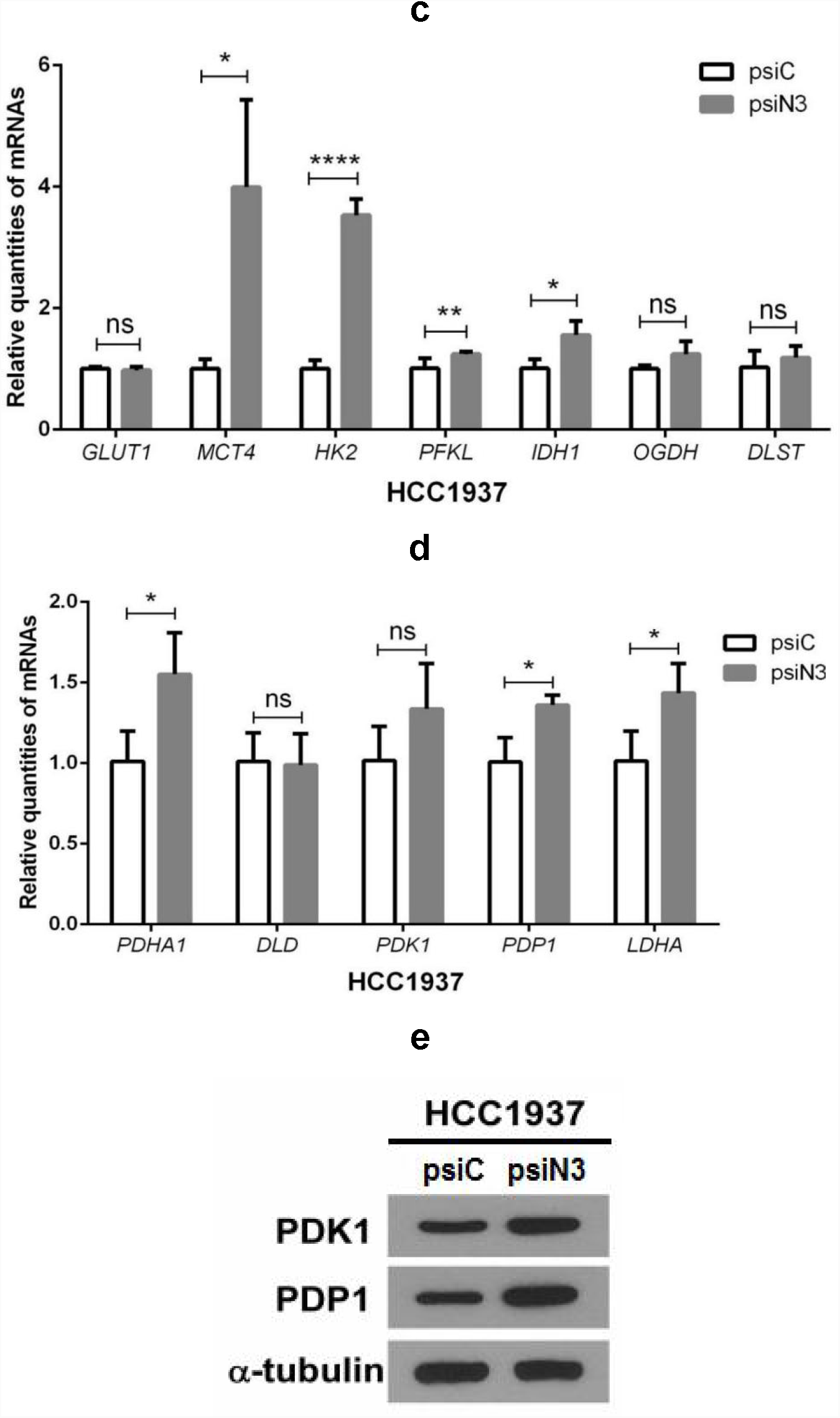
Impact of *NOS2* knockdown on glycolytic intermediate transport and aerobic/anaerobic glucose degradation in HCC1937::psiC/psiN3 mice. **a** ATP levels; **b** LA levels; **c** *GLUT1, MCT4, HK2, PFKL, IDH1, OGDH*, and *DLST* mRNA levels; **d** *PDHA1, DLD, PDK1, PDP1*, and *LDHA* mRNA levels; **e** PDK1 and PDP1 levels.

To further determine how anaerobic catabolism could be converted to aerobic catabolism, we quantified the levels of mRNAs encoding the critical substrate transporter and Krebs cycle enzymes in HCC1937::psiC/psiN3 cells. As illustrated in Fig. 3c, the mRNA levels of glucose transporter 1 *(GLUT1)* responsible for glucose intake into cells was nearly unchanged, whereas that of monocarboxylate transporter 4 *(MCT4)* responsible for the transport of pyruvate, an intermediate of aerobic glucose catabolism, into mitochondria was markedly increased, suggesting the complete degradation of glucose via pyruvate in mitochondria. Accordingly, the mRNA of hexokinase 2 *(HK2)* mRNA, which encodes an enzyme that degrades glucose to pyruvate, and 6-phosphofructokinase liver type (*PFKL*) and isocitrate dehydrogenase 1 *(IDH1)*, which encode the enzymes that degrade pyruvate, were also significantly increased; however, the mRNAs of the other enzyme-encoded oxoglutarate dehydrogenase (*OGDH)* and dihydrolipoamide S-succinyltransferase *(DLST)* were upregulated without statistical significance (Fig. 3c).

In the pivotal controlling centre determining glycolysis or Krebs cycle reactions, the pyruvate dehydrogenase complex (PDC), pyruvate dehydrogenase alpha 1 (PDHA1)-encoded mRNA was significantly elevated, although dihydrolipoyl dehydrogenase (DLD)-encoded mRNA remained unchanged. While the mRNA encoding pyruvate dehydrogenase kinase isozyme 1 (PDK1) for PDH inactivation by phosphorylation was increased without statistical significance, the mRNA encoding pyruvate hydrogenase phosphatase 1 (PDP1) for PDH activation by dephosphorylation was increased with statistical significance (P<0.05). Nevertheless, lactic acid dehydrogenase A (LDHA) was still upregulated, suggesting that the conversion from glucose to LA via glycolysis was partially maintained (Fig. 3d). As evidence confirming the co-existence of anaerobic and aerobic glucose catabolism, both PDK1 that inactivates PDH and PDP1 that activates PDH were synchronously increased (Fig. 3e).

### Enhanced AMPK/SIRT1/PGC-1α signalling cascades drive mitochondrial biogenesis but unchange angiogenesis

To evaluate the effect of *NOS2* knockdown on mitochondrial biogenesis, we quantified the expression levels of AMPK/SIRT1/PGC-1α signal transducers in HCC1937::psiC/psiN3 cells. As a result, *AMPK* (p<0.05), *SIRT1* (p<0.05), *PGC-lα* (p<0.01) and *COX4* mRNAs (p<0.05) were synchronously increased with statistical significance (Fig. 4a), although AMPK, pAMPK, Akt, pAkt, SIRT1, PGC-1α, and COX4 were only slightly increased (Fig. 4c).

**Fig. 4.**
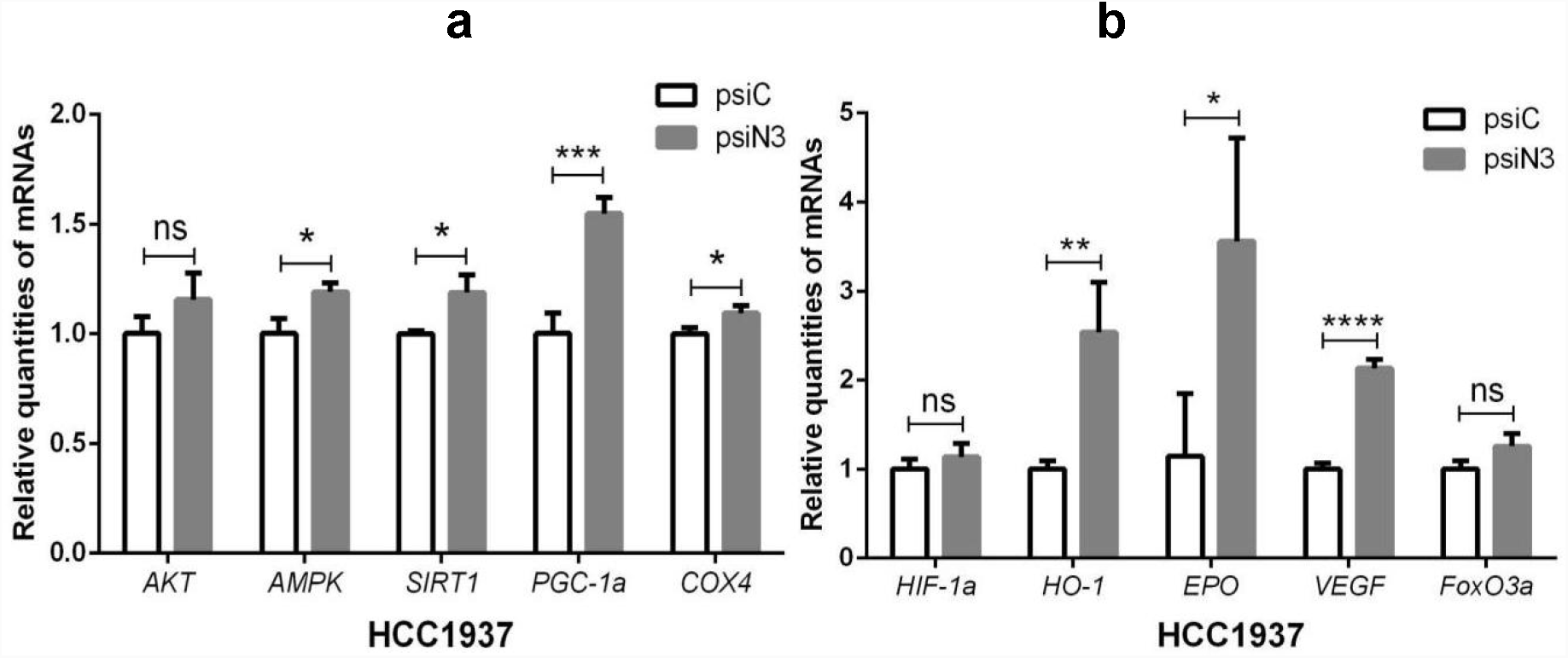
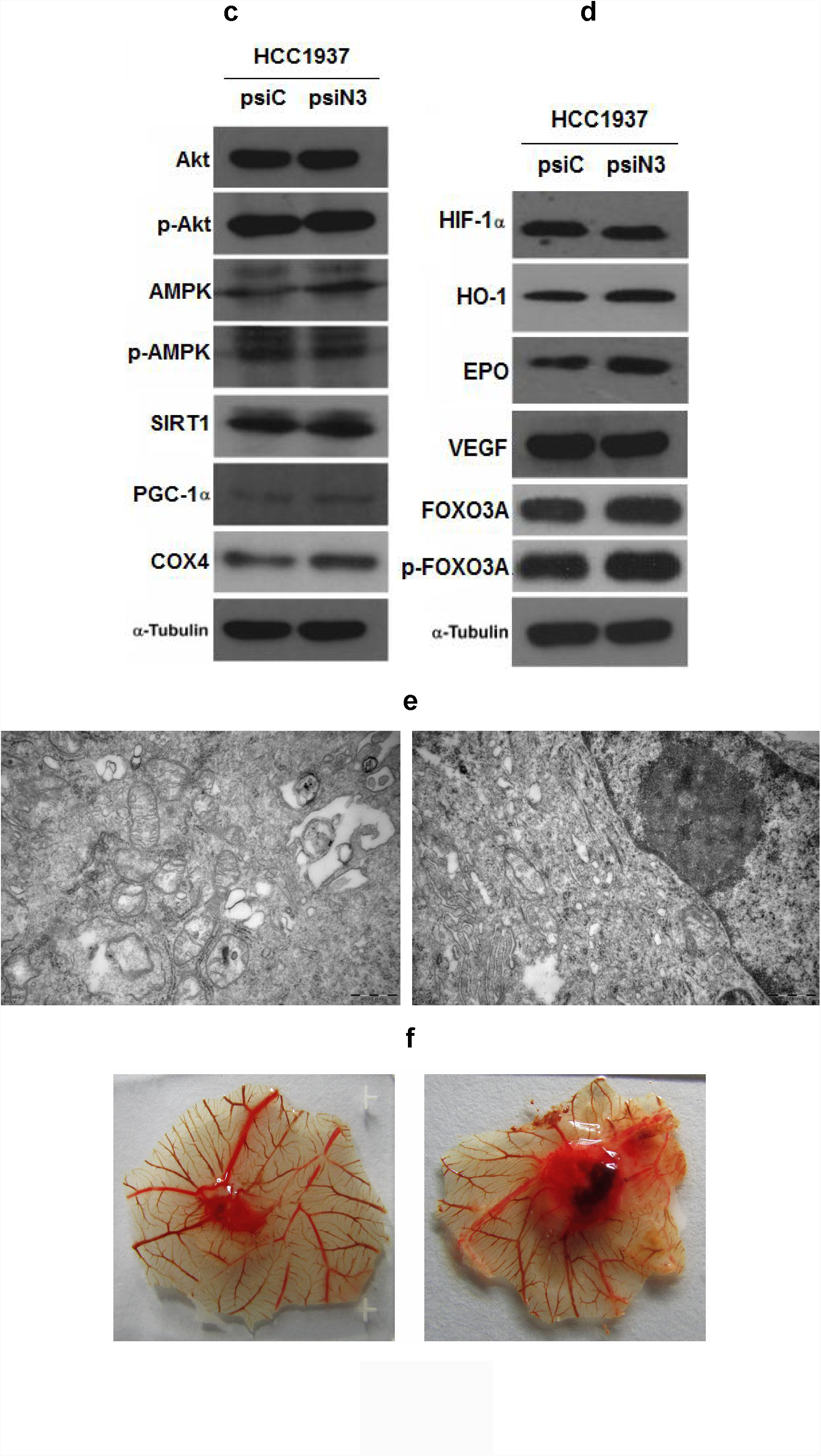
Effects of *NOS2* knockdown on the AMPK signalling pathway, mitochondrial biogenesis, and angiogenesis in HCC1937::psiC/psiN3 cells. **a** *AKT, AMPK, SIRT1, PGC-1α*, and *COX4* mRNA levels; **b** *HIF-1α, HO-1, EPO, VEGF*, and *FoxO3a* mRNA; **c** AKT, p-AKT, AMPK, p-AMPK, SIRT1, PGC-1α, and COX4 levels; **d** HIF-1α, HO-1, EPO, VEGF, and FOXO3A levels. **e** The morphology of mitochondria (50,000X) in HCC1937::psiC (left) and in HCC1937:: psiN3 cells (right). f The chick chorioallantoic membrane assay in HcCl937::psiC (left) and in HCC1937::psiN3 cells (right).

To explore whether *NOS2* knockdown might affect hypoxia-inducible genes, we quantified the expression levels of HIF-1α and some downstream genes in HCC1937::psiC/psiN3 cells. Excluding *HIF-1α* mRNA, *HO-1* mRNA(p<0.01) for anti-oxidation, *EPO* mRNA (p<0.05) for erythrogenesis, and *VEGF* mRNA (p<0.05) for angiogenesis were overexpressed upon *NOS2* knockdown (Fig. 4b). However, HIF-1α maintained a constant level despite the elevation of HO-1, EPO and VEGF (Fig. 4d), implying that HIF-1α was subjected to post-translational regulation. While the tumour suppressor-encoding *FoxO3a* mRNA increased insignificantly, FOXO3A and p-FOXO3A are increased with discernible differences.

As shown, the mitochondria were larger but scarce in HCC1937::psiC cells, whereas they were smaller but extensively distributed in HCC1937::psiN3 cells (Fig. 4e), indicating that *NOS2* knockdown might have driven mitochondrial biogenesis by activating AMPK/SIRT1/PGC-1α signalling. Additionally, no distinguishable difference in angiogenesis phenotypes was found in chick chorioallantoic membrane treated with HCC1937::psiN3 cells compared with HCC1937::psiC cells, albeit the presence of more blood knots following the treatment with HCC1937::psiN3 cells (Fig. 4f).

### *NOS2* knockdown suppresses tumour cell proliferation *in vitro* and retards tumour cell growth in nude mice

To examine whether *NOS* knockdown would repress TNBC cell proliferation, we compared the growth rates between HCC1937:: psiN3 and HCC1937:: psiC cells. After culturing *in vitro* for 5 d, a gradual decline in cell growth was observed for HCC1937::psiN3 cells. As shown in Fig. 5a, the growth rate of HCC1937::psiN3 cells was much lower than that of HCC1937::psiC cells, accounting for 35% of the decline in the absorbance at 490 nm.

**Fig. 5.**
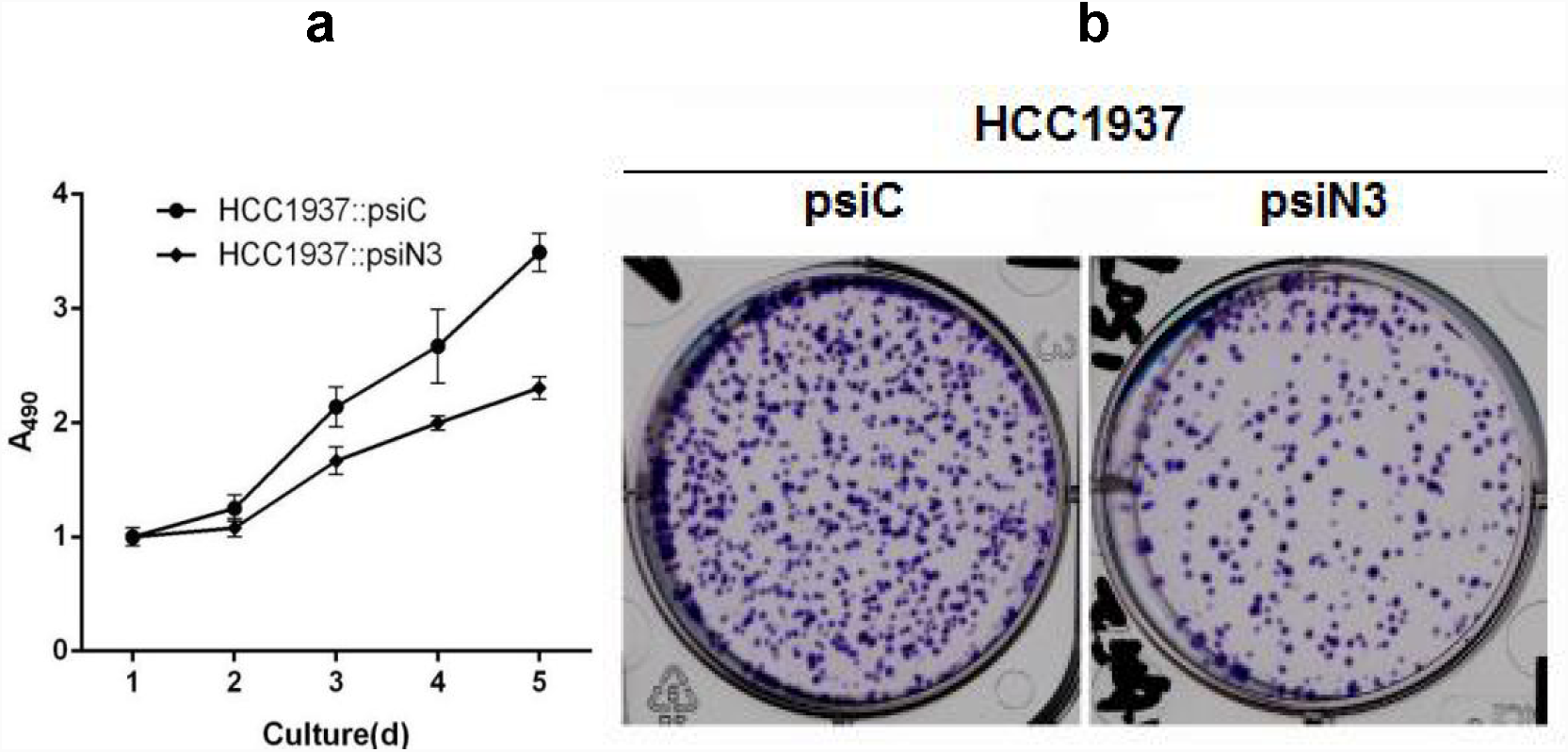
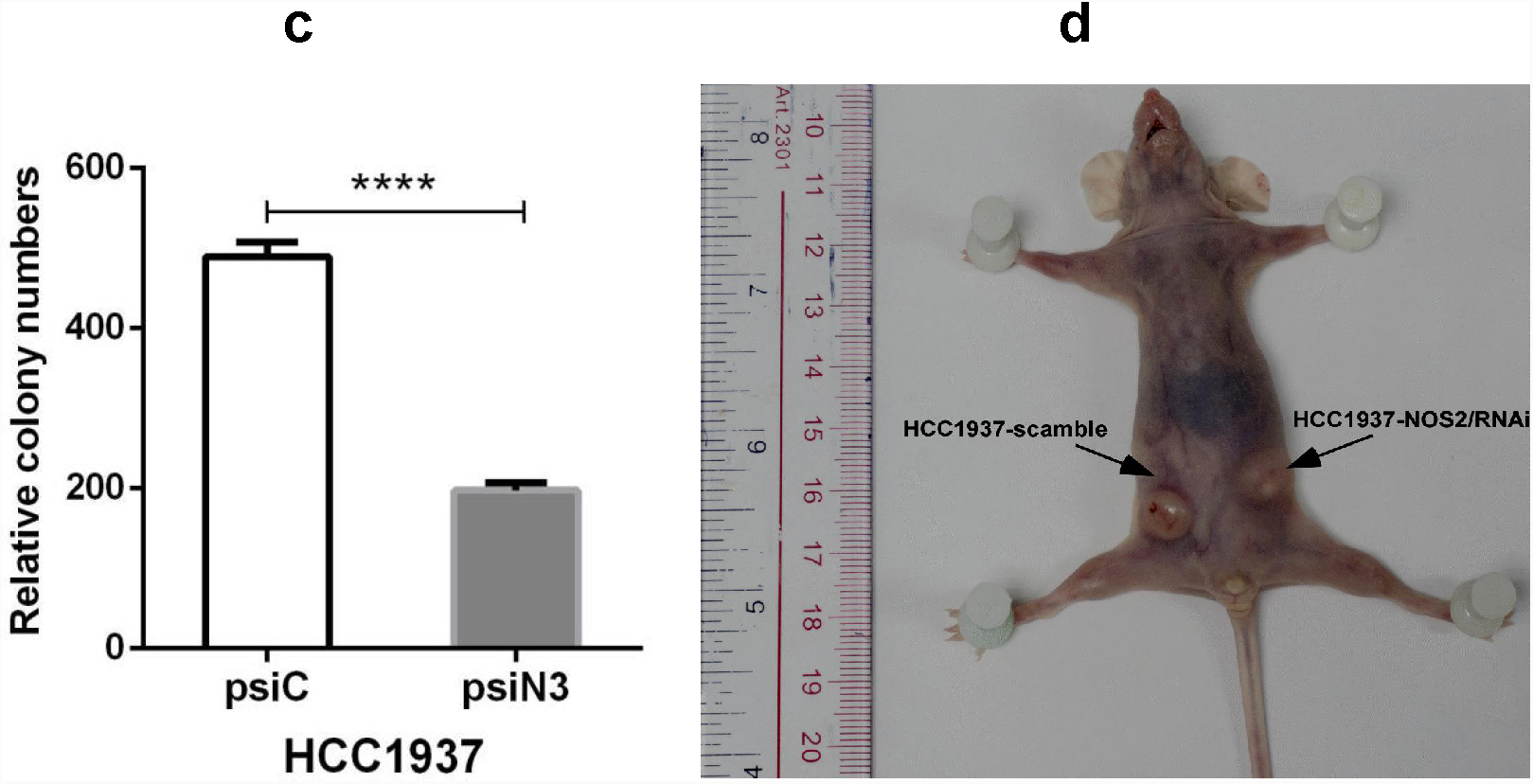
Effects of *NOS2* knockdown on cell proliferation *in vitro* and cell xenograft growth in a nude mouse. **a** the relative growth curve of HCC1937::psiC/psiN3 cells. **b** culture plates showing the colony numbers of HCC1937::psiC cells (left) and HCC1937::psiN3 cells (right). **c** the relative colony numbers of HCC1937::psiC/psiN3 cells. **d** the morphology of nude mice carrying the xenograft HCC1937::psiC/psiN3 cells.

After culturing *in vitro* for 7-10 d, fewer colonies were counted on the plate of cultured HCC1937:: psiN3 cells than HCC1937:: psiC cells (Fig. 5b). The relative colony numbers were much lower for HCC1937:: psiN3 cells than HCC1937:: psiC cells (Fig. 5c). These results supported a correlation of repressed tumour proliferation with downregulated *NOS2* expression.

In the xenograft tumour-carrying nude mice, proliferation was observed for HCC1937::psiN3 cells, but not for HCC1937::psiC cells, suggesting that tumourigenicity was partially reversed upon *NOS2* knockdown (Fig. 5d).

### *ESR1* **knockdown decreases ERα and activates cell proliferative signalling with modulation of oncogenes and tumour suppressor genes**

To silence *ESR1* and mimic ERα inactivation, three recombinant *ESR1*-targeted siRNA expression constructs, psiRNA-ER1/2/3 (abbr. psiE1/2/3), were constructed by inserting one of three candidate shRNA fragments (the sequences were aligned in Methods), *ESR1* shRNA1/2/3, into the integration-type plasmid expression vector pSUPER-retro-puro.

After separately transferring those vehicles into MCF-10A cells, *ESR1* mRNA and ERα levels were quantified in MCF-10A::psiE1/2/3 cells transiently expressing *ESR1* shRNA. It was noted that *ESR1* mRNA and ERα levels declined after transient infection of psiE1/2/3 into HCF-10A cells, as shown in Fig. 6a and 6b. From the expression levels, MCF-10A:: psiE3 cells showed the lowest *ESR1* mRNA and lowest ERα levels. Thus, psiE3 was chosen for infection of MCF-10A cells to obtain stable *ESR1* shRNA expression.

**Fig. 6.**
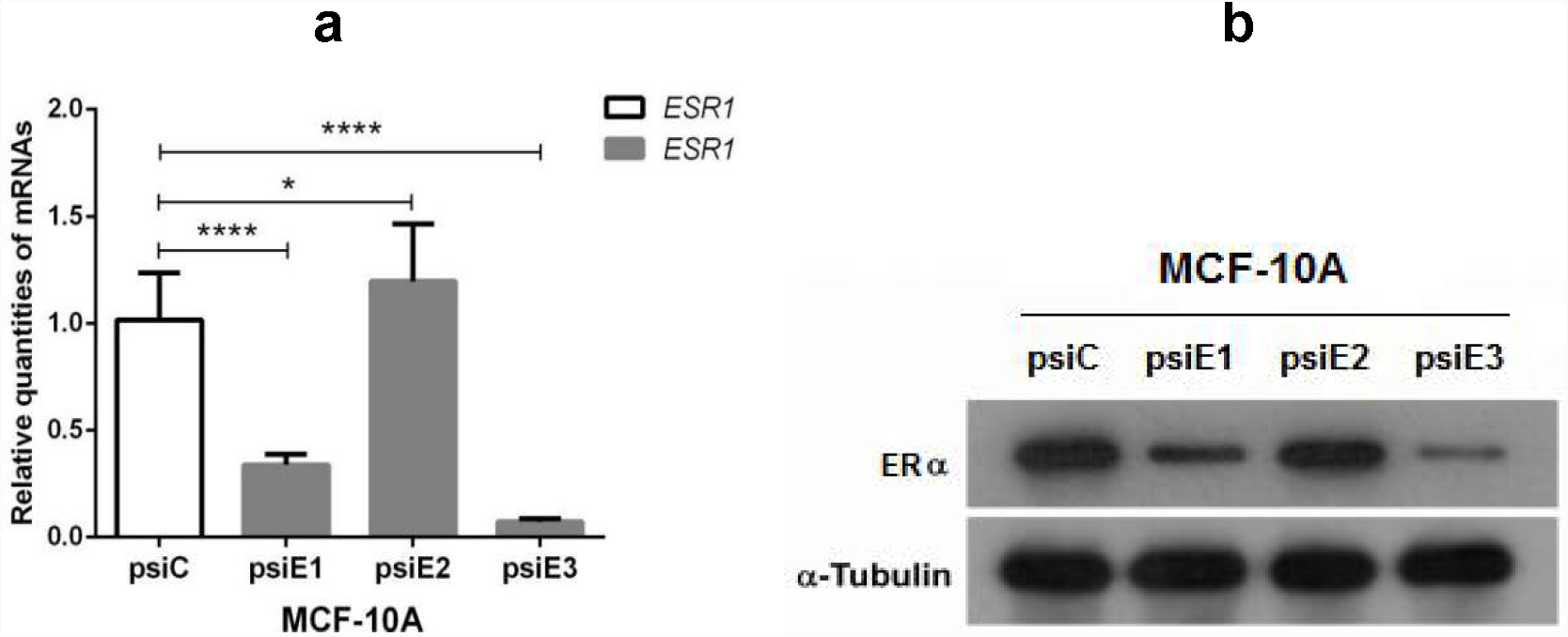
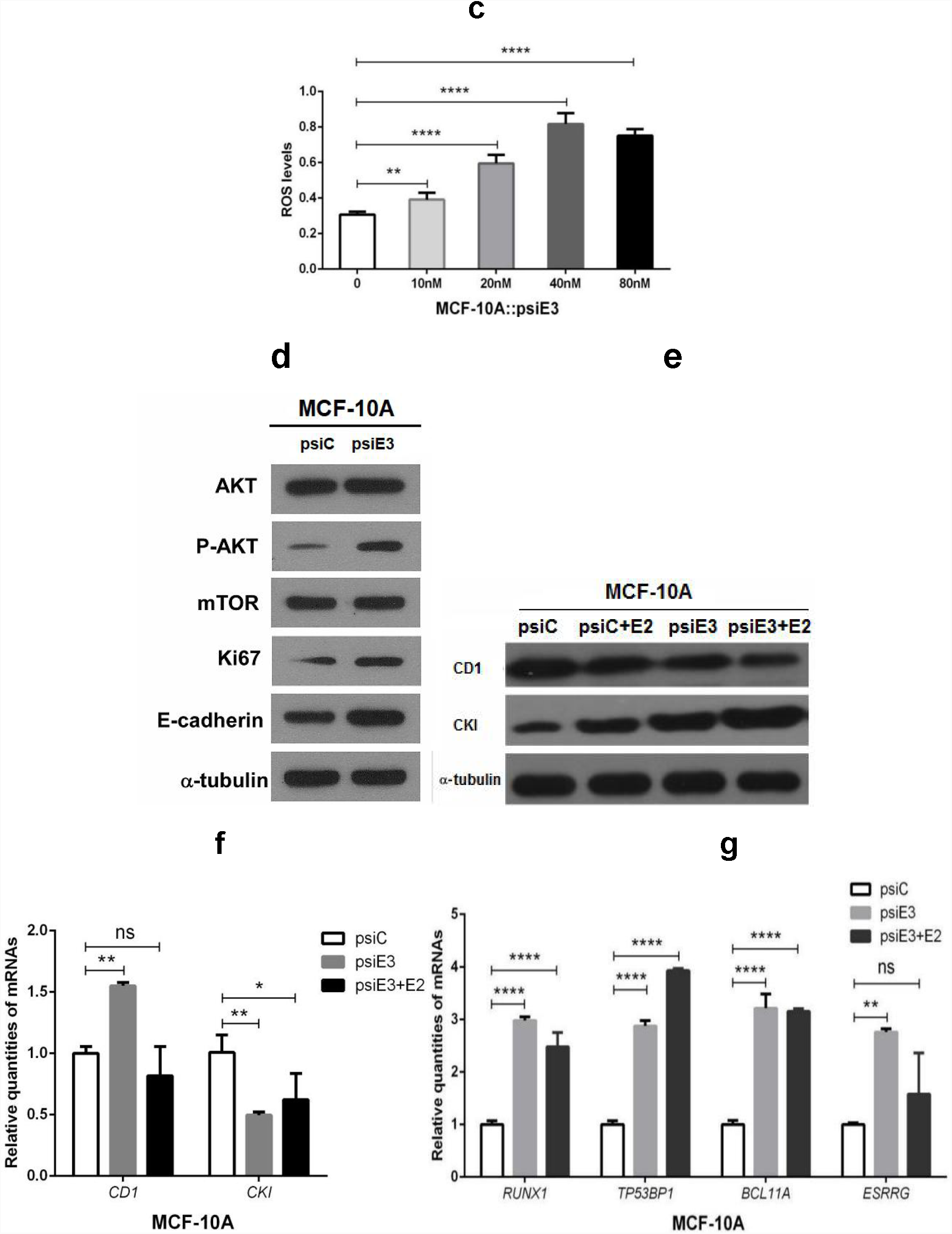
The expression profiles of cell proliferative signalling transducer genes, oncogenes and tumour suppressor genes after shRNA-guided *ESR1* knockdown in MCF-10A::psiC/psiE3 cells with or without E2. **a** *ESR1* mRNA levels (transient expression); **b** ERα levels (transient expression); **c** ROS levels; **d** Akt, p-Akt, mTOR, Ki67 and E-cadherin levels; **e** CD1 and CKI levels; **f** *CD1* and *CKI* mRNA levels; **g** *RUNX1, TP53BP1, BCL11A* and *ESRRG* mRNA levels.

To create a milieu similar to oestrogenemia due to ERα dysfunction, MCF-10A::psiC/psiE3 cells were incubated with different E2 levels (10, 20, 40, and 80 nM). After culturing for 24 h, ROS levels were measured separately in the tested cells. As shown in Fig. 6c, the ROS burst occurred in an E2 concentration-dependent manner, among which 40 nM E2 gave rise to the highest ROS levels. Thus, 40 nM E2 was chosen for the subsequent incubation experiments.

To validate how can *ESR1* knockdown cause tumourigenic changes, the expression levels of some cell proliferative signalling transducer genes, oncogenes and tumour suppressor genes were quantified in MCF-10A::psiC/psiE3 cells. As results, *ESR1* knockdown from MCF-10A cells could activate the cell proliferation-responsible PI3K/Akt/mTOR signalling pathway, in which Akt and p-Akt were upregulated but mTOR was unchanged. Interestingly, tumourigenic Ki67 and tumour suppressive E-cadherin were also synchronously upregulated, implying that tumourigenic and anti-tumourigenic roles coexisted in MCF-10A::psiE3 cells (Fig. 6d).

For other tumour markers and tumour suppressors, CD1 levels were higher in MCF-10A::psiE3 cells than in MCF-10A::psiE3 cells + E2, whereas the tumour suppressive CKI levels were lower in MCF-10A::psiE3 cells than in MCF-10A::psiE3 cells + E2 (Fig. 6e). Similarly, *CD1* mRNA levels increased in MCF-10A::psiE3 cells, whereas *CKI* mRNA levels decreased in MCF-10A::psiE3 cells + E2 (Fig. 6f).

Though specific expression in TNBC, both *RUNX1* and *BCL11A* mRNA levels were dramatically upregulated in MCF-10A::psiE3 cells, highlighting a typical feature of TNBC pathogenesis. Similarly, *TP53BP1*, which is underexpressed in most cases of TNBC, was also upregulated as a sign of DNA damage, suggesting an increase in DNA repair activity. In addition, ESRRG-encoding ERRy was upregulated with unknown functions (Fig. 6g).

These results clearly indicated that normal cells with *ESR1* knockdown experienced a tumour-like change, such as an upregulation of oncogenes and a downregulation of tumour suppressor genes, in contrast to the pattern observed in tumour cells with *NOS2* knockdown.

### *ESR1* **knockdown enhances hypoxia, represses anti-oxidation and unalters mitochondrial biogenesis and metabolism**

To evaluate whether *ESR2* knockdown would affect pro-inflammatory signaling, some pro-inflammatory cytokine and transcription factor mRNA levels were determined. Interestingly, *ESR2* knockdown was shown to strengthen pro-inflammatory signalling in mammary epithelial cells, although this process did not involve typical immune cells. As illustrated in Fig. 7a, *IL-6* mRNA and *STAT3* mRNA levels were elevated in MCF-10A::psiE3 cells but declined in MCF-10A::psiE3 cells + E2, underscoring that *ESR1* knockdown augmented pro-inflammatory signalling, whereas E2 treatment reversed the pro-inflammatory response and exerted an anti-inflammatory role.

**Fig. 7.**
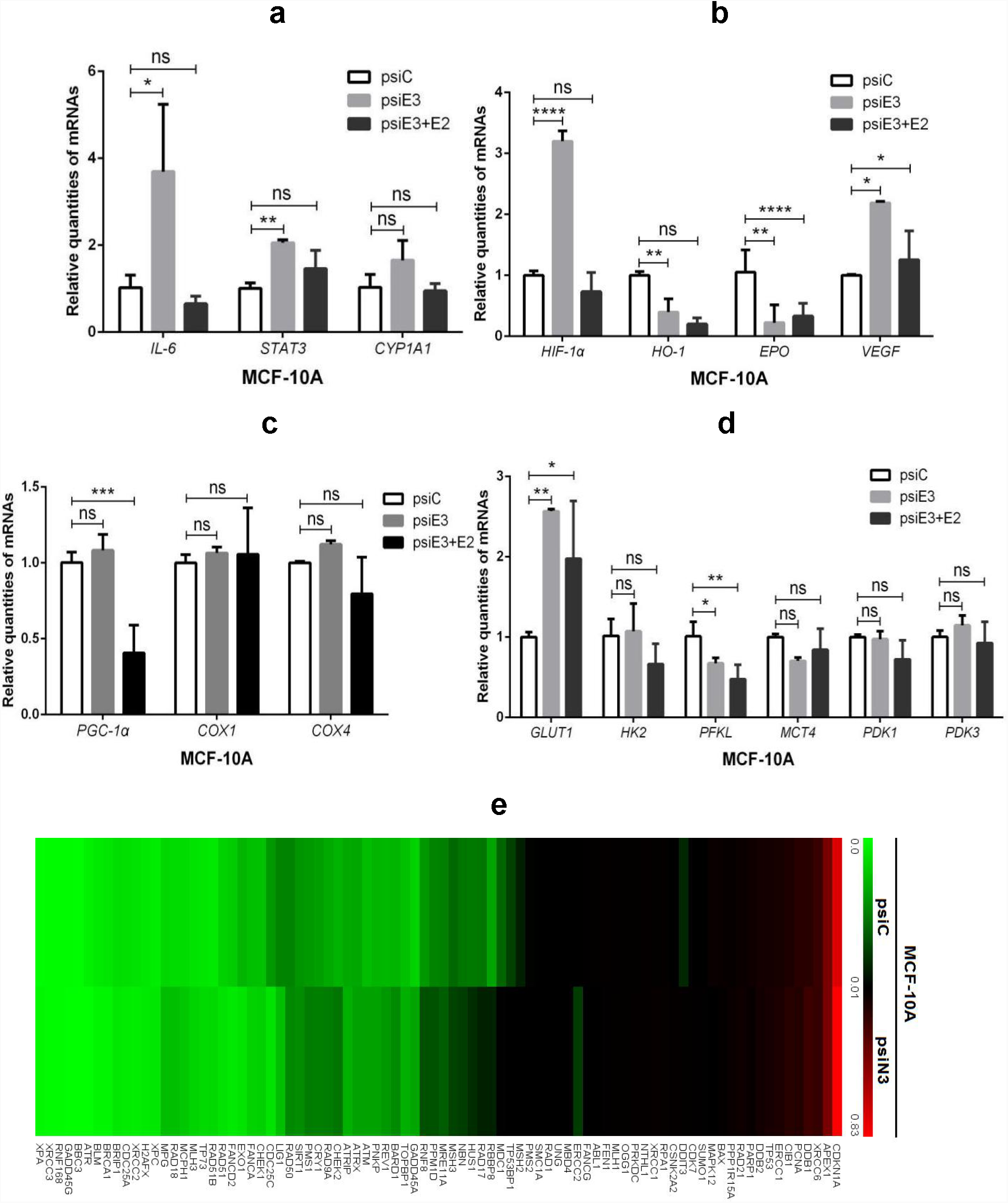
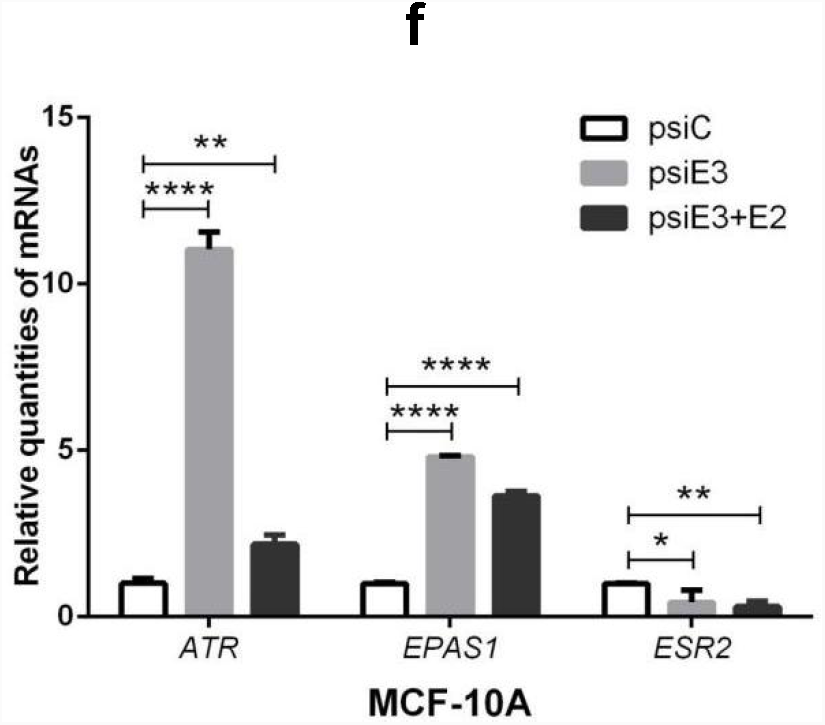
The pro-inflammatory, hypoxic and oxidative responses in MCF-10A::psiC/psiE3 cells. a *IL-6, STAT3*, and *CYP1A1* mRNA levels; b *HIF-1α, HO-1*, *EPO*, and *VEGF* mRNA levels; c *PGC-1α, COX1*, and *COX4* mRNA levels; d *GLUT1, HK2, MCT4, PDK1*, and *PDK3* mRNA levels; e DNA damage-involving mRNA levels (microarray). f *ATR, EPAS1, ESR2* mRNA levels.

Concomitantly, the upregulation of *HIF-1α* reflecting hypoxia and *VEGF* conferring angiogenesis, as well as the downregulation of *HO-1* and *EPO* endowing anti-oxidative abilities, were observed in MCF-10A::psiE3 cells, implying an induction of *HIF-1α* and *VEGF* via pro-inflammation and a repression of *HO-1* and *EPO* by anti-inflammation. In contrast, a remarkable reversion of *HIF-1α* and *VEGF* overexpression was predominantly noticed in MCF-10A::psiE3 cells + E2 (Fig. 7b).

Accordingly, the mitochondrial biomarkers encoding *PGC-1α, COX1* and *COX4* mRNA were unchanged (Fig. 7c). *GLUT1* mRNA responsible for intracellular glucose transport was upregulated in MCF-10A::psiE3 cells but downregulated in MCF-10A::psiE3+E2 cells. In contrast, *MCT4* mRNA responsible for mitochondrial pyruvate transport was downregulated in MCF-10A::psiE3 cells but upregulated in MCF-10A::psiE3+E2 cells (Fig. 7d). These results indicated that *ESR1* knockdown enhanced glycolysis in the cytosol but attenuated Krebs cycle reactions in mitochondria, although E2 could reverse those effects.

To investigate whether ERα downregulation would aggravate the inflammation-induced DNA damage, we evaluated the transcriptional profiles of 84 DNA damage pathway genes in MCF-10A::psiC/psiE3 cells by RT-PCR microarray. Consequently, DNA repairing-controlled *ATR* was upregulated 37.20-fold in MCF-10A::psiE3 cells compared with MCF-10A::psiC cells. Other genes involved in DNA damage and repair that were upregulated more than two-fold are illustrated in this clustering diagram in detail. The numbers of high-level mRNAs (red) were clearly higher in MCF-10A::psiE3 than in MCF-10A::psiC cells (Fig. 7e).

As verified by qPCR, an elevation of *ART* mRNA levels up to 10-folds was observed in MCF-10A::psiE3 cells compared with MCF-10A::psiC cells. Conversely, E2 treatment caused a significant decline in *ART* mRNA levels. Additionally, *EPAS1* mRNA encoding HIF-2α exhibited a similar pattern as *ART* mRNA, whereas *ESR2* mRNA encoding ERβ displayed a contrasting pattern to *ART* mRNA in MCF-10A::psiE3 cells, implying that the upregulation of HIF-2α was changed similarly to HIF-1α, but ERβ without upregulation did not compensate for the deficiency of ERα after *ESR1* knockdown.

### **Cells with** *ESR1* **knockdown neither displays proliferative features** *in vitro* **nor expresses tumourigenic phenotypes** *in vivo*

According to our expectations, MCF-10A::psiE3 cells should grow faster than MCF-10A::psiC cells because the former exhibited tumourigenic features after *ERS1* knockdown. However, *ERS1* knockdown cells, either with or without addition of E2, showed fewer colonies (Fig. 8a), and MCF-10A::psiE3 cells with addition of E2 showed the least relative colony numbers among the compared groups (Fig. 8b). Regarding the MCF-10A::psiE3 cells, no tumour was seen on cell grafting nude mice, demonstrating that *ESR1* knockdown was not sufficient to confer an unlimited proliferative capacity to the grafted cells (Fig. 8c).

**Fig. 8.**
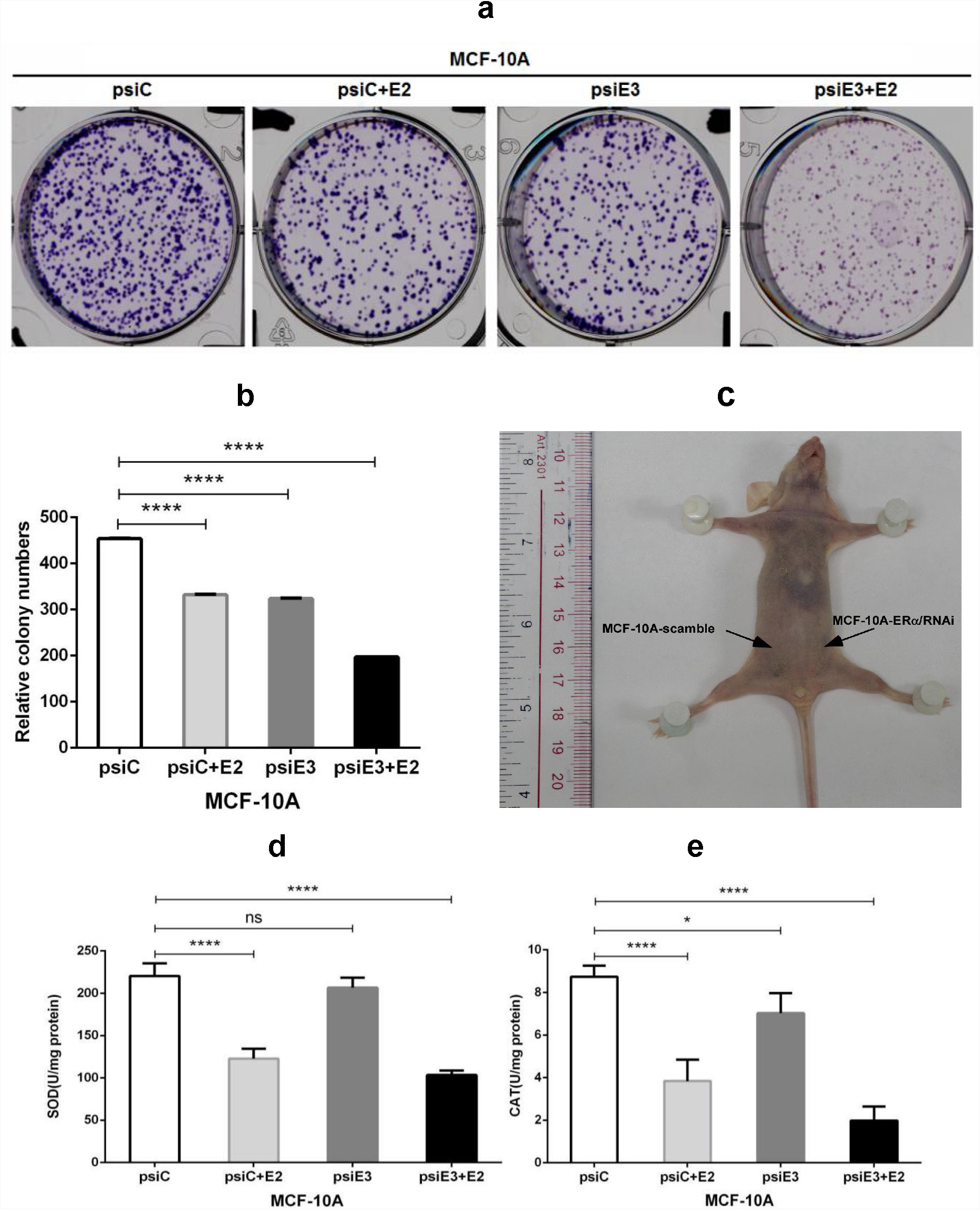
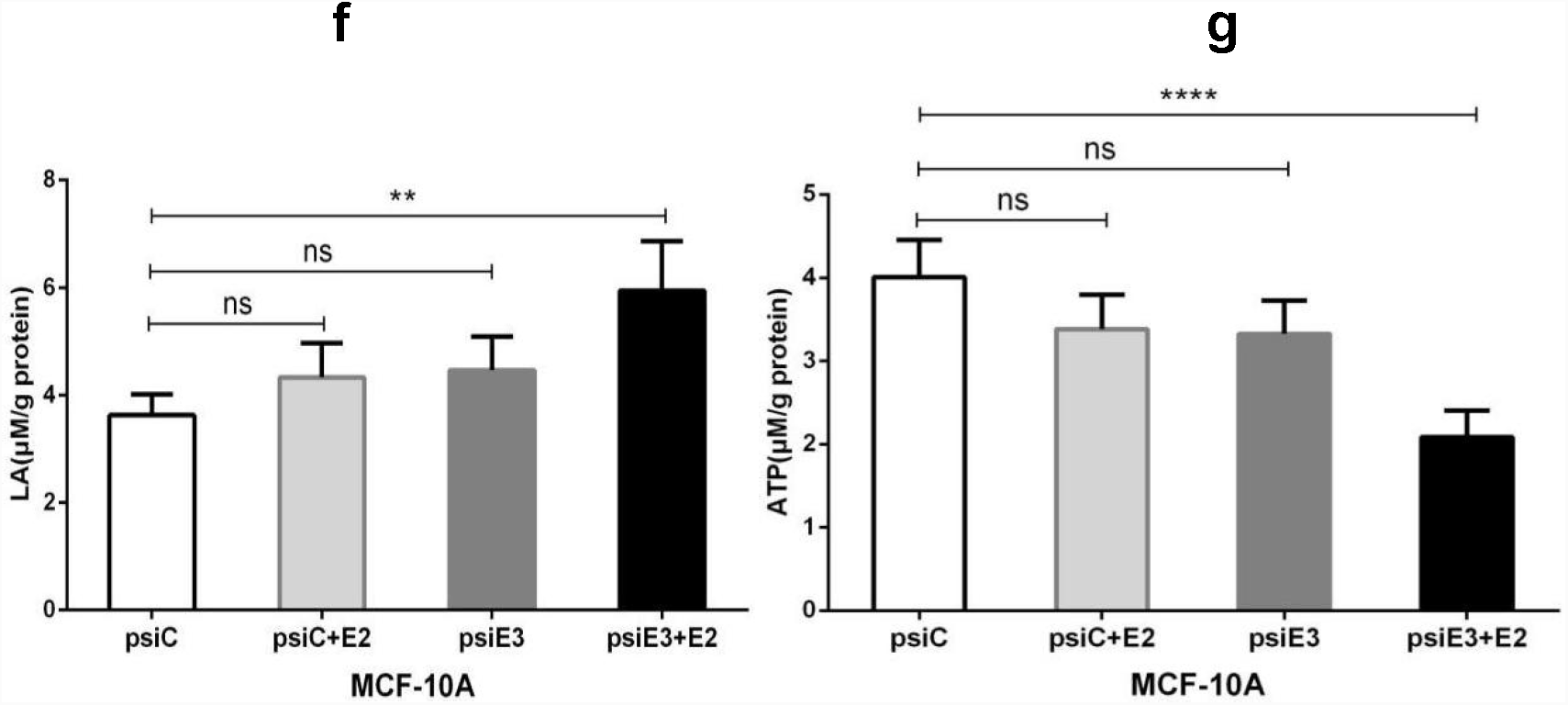
Effects of ESR1 knockdown on cell proliferation *in vitro* and cell xenograft growth on the nude mouse. **a** culture plates showing colony numbers of MCF-10A::psiC, MCF-10A::psiC+E2, MCF-10A::psiE3, and MCF-10A::psiE3+E2 cells (from the left to the right). **b** the relative colony numbers of MCF-10A::psiC/psiE3 cells with or without E2. **c** the morphology of a nude mouse carrying xenograft McF-10A::psiC/psiE3 cells, in which MCF-10A-scramble represents McF-10A::psiC and MCF-10A-ERα/RNAi represents MCF-10A::psiE3; **d** SOD levels; **e** CAT levels; **f** LA levels; **g** ATP levels.

The dramatic decreases in the relative colony numbers following exposure of MCF-10A::psiC cells to E2 might be due to E2-triggered potent ROS burst and pronounced apoptosis induction. Correspondingly, E2-triggered ROS should induce SOD and CAT to scavenge ROS, which in turn downregulate SOD and CAT. Consistent with this reasoning, MCF-10A::psiC/psiE3 cells showed almost unchanged SOD and CAT levels, whereas MCF-10A::psiE3 cells + E2 exhibited much lower SOD and CAT levels (Fig. 8d and 8e).

Accordingly, the highest LA levels and the lowest ATP levels, so-called Warburg effects observed in the tumours, were measured in MCF-10A::psiE3 cells + E2 (Fig. 8f and 8g), which seemed to be an essential consequence of E2-driven oxidative stresses although ROS was eventually scavenged by antioxidant enzymes.

These results demonstrated that *ESR1* knockdown only mimicked the tumourigenic features at the molecular level. Thus, tumourigenesis might require more time to evolve from molecular aspects to its phenotypic presentation.

### *ESR1* **knockdown reduces NO levels by dually downregulating iNOS and eNOS, coupled with compromised nitrosylation and apoptosis**

To figure out why tumour metabolic patterns were not transformed to tumour phenotypes as expectation, we assumed that the ESR1-modified cells might only exert the partial effects from ERα inactivation but not from iNOS activation. In other words, iNOS-derived NO could mediate many structural and functional changes of cellular components, which should be critical for tumourigenesis and carcinogenesis.

As shown in Fig. 9a, NO levels were reduced in MCF-10A:: psiE3 cells compared with MCF-10A::psiE3 cells + E2. Based on Fig. 9b and 9c, while unchanged or lower *NOS2* mRNA/iNOS and *NOS3* mRNA/eNOS levels were measured in MCF-10A::psiE3 cells, higher *NOS2* mRNA/iNOS and *NOS3* mRNA/eNOS levels were quantified in MCF-10A::psiE3 cells + E2.

**Fig. 9.**
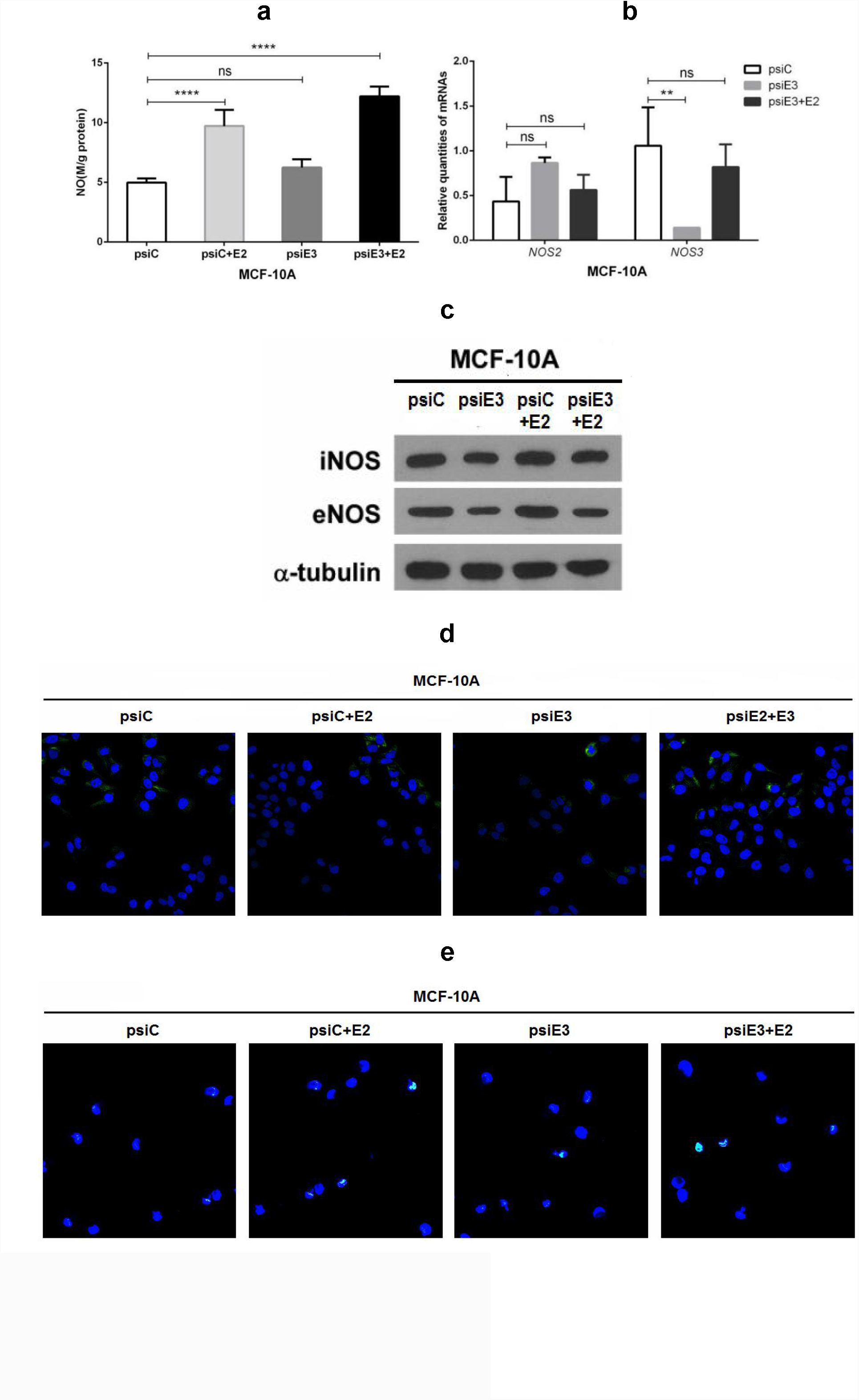
The effects of *ESR1* knockdown with or without E2 on nitrosative stress, oxidative stress, and cell death in MCF-10A::psiC/psiE3 cells. **a** NO levels; **b** *NOS2* and *NOS3* mRNA levels; **c** iNOS and eNOS levels; **d** SOD levels; **e** CAT levels; **f** nitration in MCF-10A::psiC, MCF-10A::psiC+E2, MCF-10A::psiE3, MCF-10A:: psiE3+E3 cells (from the left panel to the right panel); **g** apoptosis in MCF-10A::psiC, MCF-10A::psiC+E2, MCF-10A::psiE3, McF-10A::psiE3+E3 cells (from the left panel to the right panel).

The lower NO levels should predispose lower NT levels because NT is a tyrosine residue modified by ONOO^-^ derived from NO and ROS reactions. As observed, MCF-10A::psiC/psiE3 cells + E2 displayed lower NT levels (Fig. 9d). Similarly, MCF-10A::psiC/psiE3 + E2 cells exhibited a reduced extent of apoptotic induction (Fig. 9e).

These results indicated that no assistance with iNOS induction, *ESR1* knockdown alone might be insufficient to allow the conversion of normal cell phenotypes into tumour cell phenotypes.

## Discussion

To validate whether inflammation-driven *NOS2/iNOS* overexpression would be a point of origin for the initiation and maintenance of TNBC, we downregulated *NOS2*/iNOS in TNBC cells by shRNA-guided knockdown. *NOS2*/iNOS downregulation resulted in NOS3/eNOS upregulation, which compromised inflammatory responses, augmented mitochondrial biogenesis, and alleviated Warburg effects by converting the metabolism from anaerobic glycolysis to aerobic oxidative phosphorylation. These results were reverse-correlated with that of the transformation of normal cells to cancer cells accompanied by a switch from oxidative phosphorylation to glycolysis [25]. As expected for the mitigation of tumourigenesis, *NOS2* knockdown not only downregulated the special TNBC biomarkers *RUNXI* and *BCLIIA* but also the tumour marker CD1, and it upregulated the tumour suppressor CKI, indicating that *NOS2* knockdown partially reversed the tumourigenic features of TNBC cells.

Similar to *NOS2* knockdown, the iNOS inhibitor N^G^-monomethyl-L-arginine monoacetate (L-NMMA) blocked the tumour-like hyperplasia in our previous observations [26]. As further evidence, the pan-NOS inhibitor L-NMMA and the selective iNOS inhibitor 1400 W have been confirmed to diminish cell proliferation, cancer stem cell self-renewal, and cell migration *in vitro* [27]. These results strongly supported the conclusion that the switch of iNOS can serve as a critical check-point towards mammary differentiation or tumourigenesis. For other tumours, iNOS overexpression can result in human colon adenoma [28]; iNOS shows enhanced KRAS-induced lung carcinogenesis, inflammation and microRNA-21 expression [29]; and glioma stem cell proliferation and tumour growth are promoted by iNOS [30].

In contrast, the sustained induction of *NOS2*/iNOS leads to a potent NO burst, which enhances 3-nitration on tyrosine residues and S-nitrosylation on cysteine residues in proteins and enzymes, thereby inactivating them, including ERα and other receptors. Evidence supporting this phenomenon suggests that NO-mediated ERα nitrosylation regulates oestrogen-dependent gene transcription [31]; E2 and protein nitrosylation are synchronously augmented in the vascular endothelium [32]; and iNOS is expressed in human breast tumours, where its presence correlates with the tumour grade in patients [33].

To mimic nitrosylation-mediated ERα dysfunction, we downregulated *ESR1/ERα* in mammary epithelial cells by shRNA-directed knockdown. The results contrasted with *NOS2* knockdown-mediated partial TNBC reversal, as *ESR1* knockdown preliminarily re-enacted mammary tumourigenesis, in which tumour-like anaerobic metabolism and proliferative signalling were examined. While *RUNX1* and *BCL11A* showed a downregulation, we also noticed an upregulation of CD1 and downregulation of CKI, together with the pronounced Warburg effects and enhanced ROS burst. These results showing a conversion from aerobic glucose catabolism to anaerobic glucose catabolism coincided with the classic metabolic features of tumour cells [25].

The high levels of oestrogens were found to upregulate ER expression, which initiates HIF-1α-mediated hyperplasia involving VEGF upregulation [34]. Under healthy conditions, oestrogens can reduce lipopolysaccharide (LPS)-triggered inflammation [35] by modulating the inflammatory p38 mitogen-activated protein kinase (MAPK)/nuclear factor κB (NF-κB) cascade [36]. E2-mediated activation of serum-glucocorticoid regulated kinase 1 (SGK1) is known to suppress LPS-mediated apoptosis and promote anti-inflammatory TH2 responses in decidual stromal cells [37]. Several studies have also shown that oestrogens can alleviate oxidative stress [38). The beneficial effects of oestrogens are also manifested in the longer lifespan of women than men [39].

Regarding metabolic patterns, *NOS2* knockdown in tumour cells maximised the Warburg effect, whereas *ESR1* knockdown in normal cells minimised it, implying that breast cancer could be reversed or re-enacted by a single gene knockdown. Importantly, *NOS2*/iNOS knockdown could be simulated by iNOS inhibition through the use of 1400 W or L-NMMA, which should provide a cheap and convenient procedure for the treatment of clinically intractable TNBS patients. Considering the effectiveness of L-NMMA in decreasing tumour growth and enhancing the survival rate in TNBC, some authors have proposed to initiate a targeted therapeutic clinical trial of L-NMMA [27].

MCF-10A::psiE3 cells, either as *in vitro* cultures or *in vivo* xenografts, were not observed to proliferate as intrinsic tumour cells. This unexpected result requires further validation, both technical and mechanistical. Nevertheless, there are some hints for deciphering the *ESRR* overexpression-induced inhibition of cell proliferation [40]. Alternatively, a mechanistic link from the unchanged production of NO and ROS to the non-proliferative growth would be also established in MCF-10A cells only with *ESR1* knockdown because iNOS could not be activated under a non-inflammatory niche. Furthermore, no mutations of two tumor suppressor genes, *BRCA2* and *TP53*, were detected (unpublished data), reflecting an anti-tumor activity in MCF-10A::psiE3 cells.

Importantly, we observed the upregulation of proliferative PI3K/Akt/mTOR signalling transducers, including Akt, pAkt and mTOR, implied that tumourigenesis might have been initiated and would also be progress. There is a correlation of the specific proliferation of breast cancer with the overexpression of the cell cycle-regulated protein Ki67 [41], and thus our results reinforce that Ki67 overexpression is relevant to MCF-10A tumourigenesis. In contrast, we examined the unexpected upregulation of E-cadherin in MCF-10A cells with *ESR1* knockdown, which is conventionally considered to be a good prognostic marker in cancer. However, questions regarding the prognostic value of E-cadherin and β-catenin in ductal carcinomas indicate a complicated role of these molecules in breast cancer progression [42]. Therefore, the upregulation of E-cadherin might represent an essential response against tumourigenesis.

## Conclusions

Taken together, our experimental data support an association of the pro-inflammatory response with breast cancer occurrence via *NOS2* activation and ERα inactivation. While overexpression of TNBC biomarkers can be reversed in human TNBC cells by *NOS2* knockdown, they can also be re-enacted in human mammary epithelial cells by *ESR1* knockdown. These results should provide insight into the thorough exploration of the mysterious breast cancer origin and help to establish a targeted drug therapy regime for intractable TNBC.

## Acknowledgements

The authors thank Ms Wei-Xia Zeng in LangRi, Guangzhou, China for her assistance in the construction of the vectors for gene knockdown.

## Availability of data and materials

The datasets used and/or analysed during the current study are available from the corresponding author upon reasonable request.

## Ethics approval and consent to participate

All experiments were approved by The Animal Care Welfare Committee, Guangzhou University of Chinese Medicine, Guangzhou, China (No. SPF-2015009). The experimental protocol complies with the requirements of animal ethics issued in The Guide for the Care and Use of Laboratory Animals, National Institute of Health (NIH), USA. The number of animals used for each experiment was minimised as much as possible, and every effort was made to reduce the chance of pain or suffering.

